# Glucocorticoid receptor activation reorganizes Wnt/LEF1 regulatory circuitry associated with therapeutic response in B-cell acute lymphoblastic leukemia

**DOI:** 10.64898/2026.09.23.752002

**Authors:** Robert J. Mobley, Damilola Oluwalana, Gabriele Baniulyte, Minzhang Zheng, Brett Brown, Faten Usrof, Kashi Raj Bhattarai, Kelly R. Barnett, Virginia Valentine, Marc Valentine, Seth Staller, Satoshi Yoshimura, Kristine R. Crews, David T. Teachey, Steven Burden, Yong Cheng, Jun J. Yang, Daniel Savic

**Affiliations:** Department of Pharmacy and Pharmaceutical Sciences, St. Jude Children’s Research Hospital, Memphis TN 38105, USA; Hematological Malignancies Program, St. Jude Children’s Research Hospital, Memphis TN 38105, USA; Graduate School of Biomedical Sciences, St. Jude Children’s Research Hospital, Memphis, TN 38105, USA; Department of Hematology, St. Jude Children’s Research Hospital, Memphis TN 38105, USA; CytoStem Shared Resource, St. Jude Children’s Research Hospital, Memphis TN 38105, USA; Department of Pediatrics, The University of Pennsylvania Perelman School of Medicine, Philadelphia PA 19104, USA; Division of Oncology, The Children’s Hospital of Philadelphia, Philadelphia PA 19104, USA

**Author notes:** Corresponding Author, Daniel Savic.

## Abstract

Glucocorticoids (GCs) are foundational to B-cell acute lymphoblastic leukemia (B-ALL) therapy, yet how glucocorticoid receptor (GR) activation interacts with lineage regulatory circuitry to shape therapeutic response remains incompletely understood. Using LEF1 to interrogate canonical Wnt regulatory circuitry, we show that GR activation reorganizes GR+LEF1 elements into recruitment, retention, and dissociation states. LEF1 dissociates from Wnt-responsive regulatory elements while being recruited to GRE-rich GR-associated elements that progressively acquire chromatin accessibility, H3K27ac, BRD4 and BRG1. RUNX1 and AP-1 show class-dependent redistribution across LEF1-defined states, extending organization beyond LEF1. GR+LEF1 reorganization is observed in patient-derived B-ALL. GR+LEF1 elements lose accessibility in GC-resistant B-ALL and during diagnosis-to-relapse evolution, with recruitment elements enriched for remodeling. Perturbation of resistance-associated GR+LEF1 elements at BIM and BMF alters GC-induced apoptotic signaling. Together, these findings identify Wnt/LEF1 circuitry as an interface through which therapeutic GR activation reorganizes the B-ALL regulatory landscape into states associated with glucocorticoid response and resistance.

## INTRODUCTION

Glucocorticoids (GCs) are a cornerstone of contemporary multi-drug therapy for B-cell acute lymphoblastic leukemia (B-ALL) and have been integral to B-ALL treatment regimens for decades (1–3). The early response of leukemic blasts to GC therapy has substantial prognostic value, with poor prednisone response consistently associated with inferior treatment outcome (2–7). Accordingly, intrinsic or acquired GC resistance remains an important barrier to successful therapy and has motivated sustained efforts to define the molecular determinants of GC sensitivity in B-ALL (1–3).

GC activity is mediated largely through the glucocorticoid receptor (GR; *NR3C1*), a ligand-activated nuclear receptor transcription factor (TF) that translocates to the nucleus and regulates transcription through direct DNA binding at *cis*-regulatory elements and interactions with other transcriptional regulators and co-factors (8,9). In B-ALL cells, GR activation initiates a transcriptional program that promotes growth arrest and apoptosis (1–3). The apoptotic response reflects coordinated regulation of multiple pathways, including activation of BH3-only BCL2-family proteins (10,11). Thus, the therapeutic response to GCs is ultimately encoded through the interaction of activated GR with the gene-regulatory circuitry of leukemia cells.

Genome-wide studies have established that this relationship is strongly influenced by the regulatory landscape that exists before hormone exposure. Up to 95% of induced GR binding occurs within pre-existing accessible chromatin, indicating that baseline chromatin state is a major determinant of GR occupancy (12). Pre-bound transcription factors can further shape receptor recruitment; for example, AP-1 factors can maintain accessible regulatory elements and facilitate subsequent GR binding (13). These and related studies have established an important paradigm in which DNA sequence, chromatin accessibility, and pre-existing TF occupancy determine where GR engages the genome.

However, the relationship between GR and the pre-existing regulatory landscape is not necessarily unidirectional. Steroid receptor activation can reprogram FOXA1 occupancy through dynamic assisted-loading mechanisms, GR can facilitate CREB1 recruitment to fasting-responsive enhancers, and GR activation can redistribute transcriptional regulators including AP-1 factors, CEBPB, and EP300 (14–17). Studies in ALL have likewise identified distinct GR-associated nucleosome-remodeling states, locus-specific recruitment of cooperating transcriptional regulators, and GC-dependent eviction of the lineage factor PU.1 linked to therapeutic response (18–21). Together, these studies establish that GC signaling can remodel chromatin and regulatory-factor occupancy in ALL.

Our previous studies provided a biologically grounded rationale for using Wnt regulatory circuitry to investigate how GR activation impacts the B-ALL regulatory genome. We linked regulation of the Wnt regulatory cofactor TLE1 to GC resistance and treatment response in patients and subsequently identified gene-regulatory crosstalk and mutual antagonism between GC and canonical Wnt signaling in B-ALL, including changes in LEF1 and TCF7L2 occupancy following GC exposure (22,23). Complementing these observations, recent chromatin-proteomic work in B-ALL further identified multiple B-cell lineage TFs, including LEF1 and RUNX1, within the DEX-dependent GR chromatome (24), and interactions between GC and canonical Wnt signaling pathways have also been described in other model systems (25–27). Together, these observations suggested that Wnt regulatory circuitry could provide a conceptual framework for understanding how therapeutic GR activation impacts the B-ALL regulatory genome and whether these changes are associated with GC response and resistance. LEF1, a major transcriptional effector of canonical Wnt signaling whose dysregulation has been associated with clinical outcome and relapse in B-ALL (28–31), therefore provided a mechanistically grounded entry point for resolving this interaction.

Here, using LEF1 to resolve the regulatory response to GR activation, we identify distinct recruitment, retention, and dissociation states with accompanying enhancer remodeling. We show that dissociation preferentially occurs at pre-existing Wnt-responsive LEF1 elements, whereas recruitment occurs at largely distinct GR-associated elements, revealing divergent outcomes of GR-Wnt/LEF1 regulatory crosstalk. Direct profiling of RUNX1 and AP-1 demonstrates that this organization extends beyond LEF1, and we connect these states to DNA sequence, progressive changes in chromatin accessibility, cofactor engagement, transcriptional response, and GC sensitivity. We further identify corresponding GR+LEF1 regulatory behavior in patient-derived leukemia and find that resistance-associated GR+LEF1 elements show lower accessibility in GC-resistant patient samples. In parallel, GR+LEF1 elements preferentially lose accessibility during diagnosis-to-relapse evolution, with recruitment elements enriched for remodeling. Functional perturbation of resistance-associated elements at BCL2L11/BIM and BMF links components of this regulatory architecture to glucocorticoid-induced apoptotic signaling. Together, these findings identify Wnt/LEF1 regulatory circuitry as a central interface through which therapeutic GR activation reorganizes the B-ALL regulatory landscape.

## RESULTS

### Glucocorticoid receptor activation redistributes LEF1 across the regulatory landscape

Building on our previous observation that GC treatment broadly alters LEF1 occupancy in B-ALL, we sought to define the genomic organization, kinetics, and regulatory context of this response. LEF1 CUT&RUN was performed across four B-ALL cell lines after 24 h DMSO or dexamethasone (DEX, 1 *μ*M) treatment. Among 78,029 LEF1 peaks, 28,059 showed decreased occupancy, 532 showed increased occupancy, and 49,438 remained unchanged (**Figure 1A**). Integration with GR ChIP-seq across five cell lines and six B-ALL PDXs (**Supplemental Figure 1 A-B)** identified 13,088 shared GR+LEF1 sites which separated into three classes: 456 recruitment sites gained LEF1 (REC), 9,576 retention sites maintained LEF1 (RET), and 3,056 dissociation sites lost LEF1 (DIS) (**Figure 1B-D**; **Table S1**).

**Figure 1.**
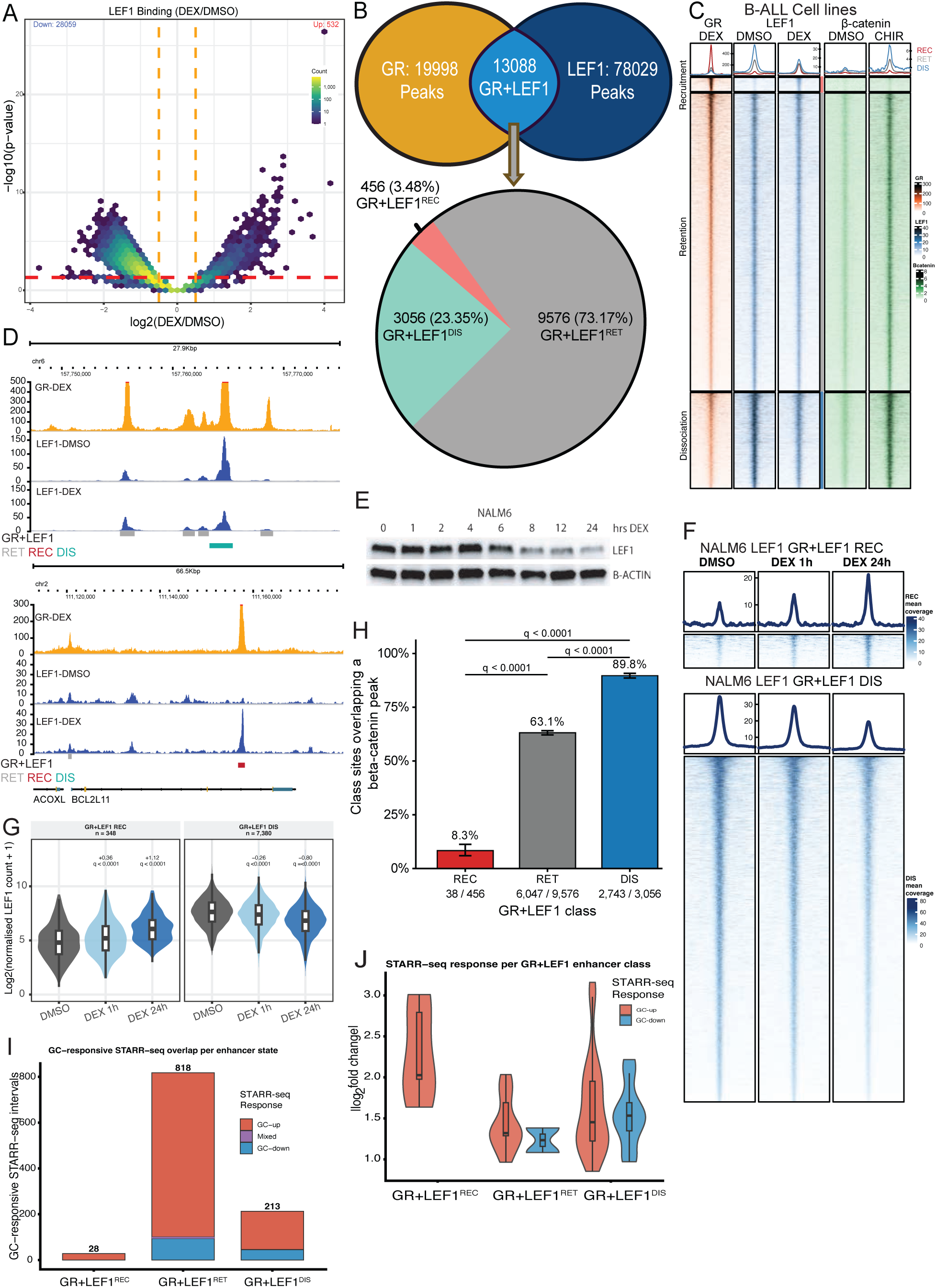
Glucocorticoid treatment defines distinct classes of GR+LEF1 regulatory interactions in B-ALL. **(A)** Volcano plot of differential LEF1 occupancy following 24 h treatment with 1 μM dexamethasone (DEX) relative to vehicle (DMSO) across 78,029 LEF1 peaks from 697, NALM6, RS411, and SUPB15 cells. DEX decreased LEF1 occupancy at 28,059 sites and increased occupancy at 532 sites, with 49,438 sites unchanged. **(B)** Overlap of GR and LEF1 binding sites following DEX treatment identifies 13,088 GR+LEF1 sites. GR binding was profiled in 5 B-ALL cell lines (697, NALM6, RS411, SUPB15 and SEM) and six PDX samples. Based on the LEF1 response to DEX, these sites were classified as recruitment (REC; n = 456), retention (RET; n = 9,576), or dissociation (DIS; n = 3,056) sites. **(C)** Heatmaps of GR, LEF1, and β-catenin occupancy across REC, RET, and DIS sites. GR and LEF1 are shown following DMSO or 24 h DEX treatment, and β-catenin following DMSO, 3 μM CHIR99021 (CHIR), or DEX+CHIR treatment. Sites are ordered by LEF1 response to DEX within each class. **(D)** Representative genomic loci illustrating REC, RET, and DIS sites and corresponding GR and LEF1 occupancy following DMSO or DEX treatment. **(E)** Western blot of LEF1 protein abundance during a DEX time course in NALM6 cells. LEF1 protein remains relatively stable through 4 h, begins to decrease by 6 h, and is substantially reduced at later time points. β-actin serves as a loading control. **(F)** Heatmaps of LEF1 occupancy at NALM6-specific recruitment (REC) and dissociation (DIS) sites following DMSO or 1 h and 24 h DEX treatment. **(G)** Distribution of LEF1 signal at NALM6-specific recruitment (REC) and dissociation (DIS) sites following DMSO or 1 h and 24 h DEX treatment. REC sites show increased LEF1 occupancy by 1 h that is further increased at 24 h, whereas DIS sites show reduced LEF1 occupancy by 1 h with a further decrease at 24 h. **(H)** Overlap of β-catenin peaks with REC, RET, and DIS sites. β-catenin occupancy is strongly enriched at DIS sites (89.8%; 2,743/3,056) and RET sites (63.1%; 6,047/9,576), but is uncommon at REC sites (8.3%; 38/456). β-catenin peaks were defined following 24 h treatment with 3 μM CHIR across the four-cell-line panel. **(I)** Overlap of previously identified glucocorticoid-responsive STARR-seq elements with REC, RET, and DIS sites. **(J)** Absolute log₂ fold change in glucocorticoid-responsive STARR-seq reporter activity across REC, RET, and DIS sites. STARR-seq data are from the previously published dataset indicated in the text.

The relationship between GR occupancy and LEF1 redistribution showed a marked asymmetry between recruitment and dissociation. LEF1 recruitment was strongly concentrated at GR-bound regions, occurring at 3.48% of GR+LEF1 versus 0.12% of LEF1-only sites (∼31-fold enrichment, p<0.0001; **Figure S1C**). In contrast, dissociation was widespread and predominantly occurred outside local GR occupancy.

Because DEX reduces LEF1 protein abundance (**Figure S1D**), we asked whether occupancy changes preceded this decrease. LEF1 protein remained largely intact through 4 h and declined thereafter, whereas redistribution was readily apparent after 1 h DEX in NALM6 cells (**Figure 1E-G**; **Figure S1D-E**). At 1 h, 14,875 peaks showed decreased and 1,248 increased occupancy; by 24 h, changes strengthened at NALM6-specific LEF1 dissociation (median log_2_ fold change - 0.26 to -0.8) and recruitment (+0.36 to +1.12) sites. Thus, LEF1 redistribution begins rapidly and precedes detectable protein depletion.

We next asked whether these response classes correspond to distinct pre-existing regulatory states. LEF1 occupancy alone does not establish engagement of canonical Wnt signaling because transcriptional activation at LEF/TCF-bound elements depends on recruitment of *β*-catenin (32,33). Following Wnt activation with CHIR99021 (CHIR, 3 *μM*), β-catenin occupied 89.8% of DIS sites, 63.1% of RET sites, and only 8.3% of REC sites (**Figure 1C,H**; **Figure S1F-G**). Thus, sites from which LEF1 dissociates following GR co-occupancy are highly enriched for pre-existing LEF1 regulatory elements capable of engaging *β*-catenin (p<0.0001), whereas GR-associated sites that recruit LEF1 are largely distinct from this Wnt-responsive state.

Finally, we examined previously generated STARR-seq data to determine whether the three classes contain GC-responsive regulatory elements (22). GC-responsive STARR-seq sites were significantly enriched at GR+LEF1 sites, with 47% overlapping this regulatory space (1,059 out of 2,269 GC-responsive sites; odds ratio=1.7, Fisher’s Exact test, p=1.18e-32). Responsive activity was identified at REC, RET, and DIS sites, while repressive activity was restricted to RET and DIS sites (**Table S2**; **Figure 1I-J**).

Together, these results reveal a rapid and asymmetric redistribution of LEF1 following GC treatment. LEF1 recruitment is highly concentrated at GR-bound regulatory elements, whereas dissociation is widespread, and begins before LEF1 protein declines; within the GR+LEF1 subset, DIS sites preferentially represent pre-existing Wnt-responsive regulatory elements. These findings define distinct recruitment, retention, and dissociation states through which GC signaling reorganizes the LEF1 regulatory network in B-ALL.

### Glucocorticoid-induced LEF1 redistribution is coupled to class-specific chromatin remodeling

GR+LEF1 sites overlapped previously published GC-responsive STARR-seq elements, supporting their capacity to function as active regulatory elements. We therefore asked whether LEF1 recruitment, retention, and dissociation were accompanied by distinct changes in chromatin state. ATAC-seq and H3K27ac ChIP-seq revealed strongly class-specific responses to DEX (**Figure 2A-B**). Among REC sites, 83.8% gained both accessibility and H3K27ac and 97.4% gained at least one of the two signals. RET sites showed a more modest, gain-biased response, whereas DIS sites were biased toward loss, with 24.4% losing at least one signal compared with 6.4% gaining. Thus, DEX-induced LEF1 redistribution is accompanied by coordinated, class-specific chromatin remodeling, with recruitment associated with strong activation of regulatory elements and dissociation associated with their reduced activity.

**Figure 2.**
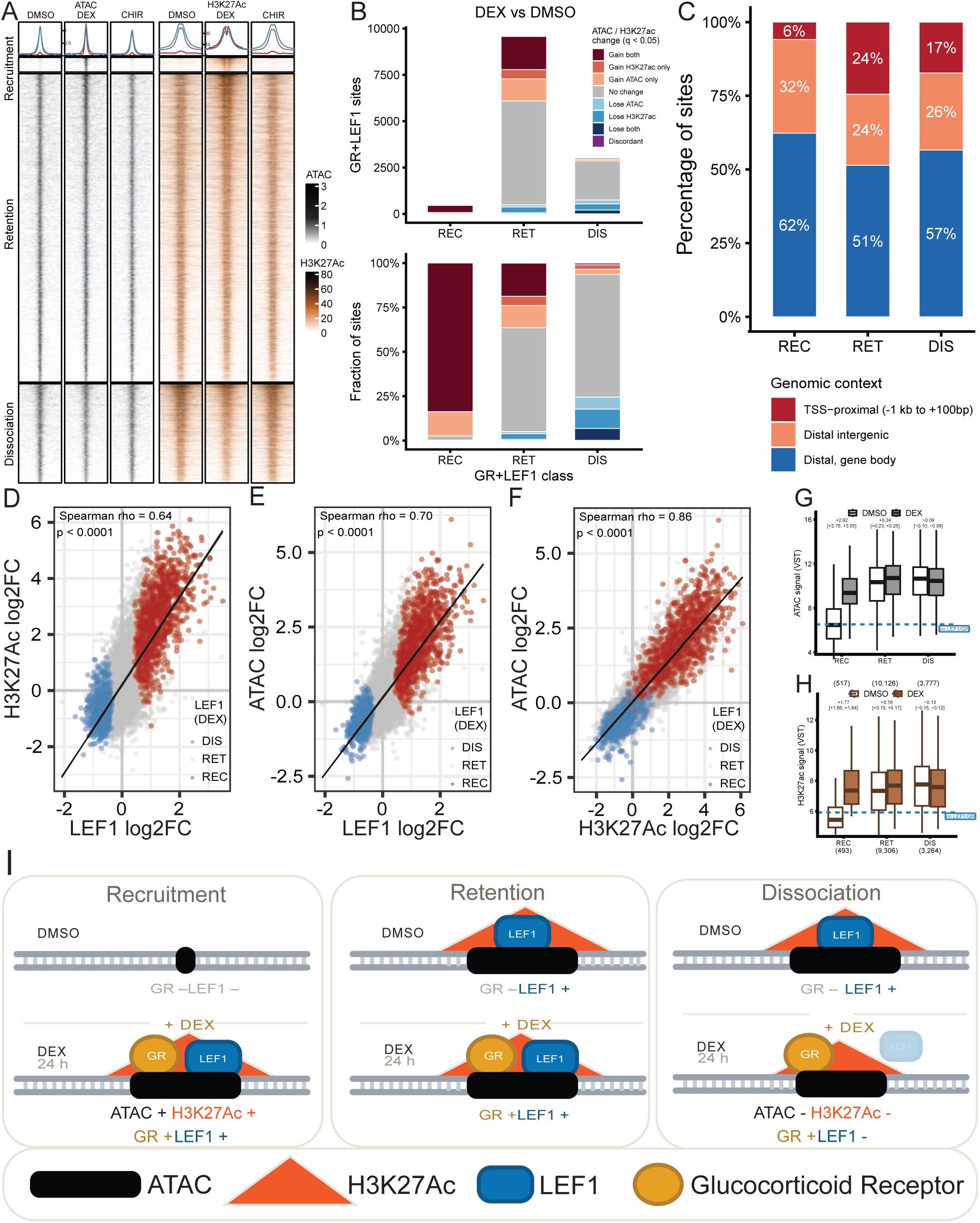
Glucocorticoid-induced LEF1 redistribution is coupled to class-specific remodeling of regulatory elements. **(A)** ATAC-seq and H3K27ac signal at REC, RET and DIS sites following vehicle (DMSO), dexamethasone (DEX), or CHIR99021 (CHIR) treatment (24 h). **(B)** Number (top) and proportion (bottom) of REC, RET, and DIS sites exhibiting significant changes in chromatin accessibility and/or H3K27ac following DEX treatment relative to DMSO (q < 0.05). Sites are classified according to significant gain or loss of ATAC-seq and H3K27ac signal, no significant change in either signal, or discordant changes between the two chromatin features. REC sites predominantly gained both accessibility and H3K27ac (83.8%; 382/456), whereas DIS sites preferentially exhibited loss of one or both chromatin features. RET sites showed an intermediate response biased toward chromatin gains. **(C)** Genomic distribution of REC, RET, and DIS sites relative to annotated genes. Sites were classified as promoter-proximal (−1 kb to +100 bp relative to the transcription start site [TSS]), intragenic, or intergenic. REC sites were less frequently promoter-proximal (5.9%) than RET (24.5%) or DIS (17.3%) sites (Pearson χ² = 142.6, df = 4, p = 7.8e-30). **(D-F)** Relationship between DEX-induced changes in LEF1 occupancy, H3K27ac, and chromatin accessibility at consensus H3K27ac peaks overlapping GR-bound sites. Scatterplots show log₂ fold changes following DEX treatment for **(D)** LEF1 versus H3K27ac, **(E)** LEF1 versus ATAC-seq, and **(F)** H3K27ac versus ATAC-seq. Spearman correlation coefficients are indicated. Sites with significantly increased or decreased LEF1 occupancy are distinguished from sites without significant LEF1 changes. **(G)** ATAC-seq signal before and after DEX treatment at REC, RET, and DIS sites. Values represent variance-stabilized signal across consensus ATAC-seq intervals overlapping each GR+LEF1 class. The dashed line indicates the median DMSO signal of reference ATAC-seq peaks lacking GR and LEF1 occupancy. REC sites exhibit low baseline accessibility followed by a marked increase after DEX, whereas changes at RET and DIS sites are substantially smaller. **(H)** H3K27ac signal before and after DEX treatment at REC, RET, and DIS sites. Values represent variance-stabilized signal across consensus H3K27ac intervals overlapping each GR+LEF1 class. The dashed line indicates the median DMSO signal of reference H3K27ac peaks lacking GR and LEF1 occupancy. REC sites exhibit low baseline H3K27ac followed by a marked increase after DEX, reaching levels comparable to untreated RET sites, whereas changes at RET and DIS sites are more modest. **(I)** Schematic showing the three GR+LEF1 classes and their responses to DEX treatment.

CHIR treatment has a comparatively limited effect at GR+LEF1 sites: 87.5% of REC sites were unchanged and only 2.4% gained both accessibility and H3K27ac, compared with 83.8% after DEX (**Figure 2A**; **Figure S2A**). RET and DIS sites likewise showed mixed responses to CHIR, indicating that Wnt activation alone did not consistently alter chromatin state across GR+LEF1 sites.

All three classes were predominantly distal to annotated transcription start sites (TSSs), but REC sites were particularly depleted from promoter-proximal regions (Figure 2C). Only 5.9% of REC sites localized within the promoter-TSS window, compared with 24.5% of RET and 17.3% of DIS sites (p=7.8e-30), indicating that GR-associated LEF1 recruitment occurs predominantly at distal elements.

To determine whether changes in LEF1 occupancy were coupled to chromatin remodeling at the same regulatory elements, we quantified LEF1, H3K27ac, and ATAC-seq signal over a shared set of consensus H3K27ac peaks overlapping GR sites (**Figure 2D-F**). Across these elements, DEX-induced changes in LEF1 occupancy correlated with changes in H3K27ac (Spearman rho=0.64) and accessibility (rho=0.70), while H3K27ac and accessibility were themselves strongly correlated (rho=0.86; all p<1.0e-4). REC and DIS sites separated in opposing directions across these comparisons. Thus, at GR-associated regulatory elements, redistribution of LEF1 is tightly coupled to the direction and magnitude of chromatin remodeling.

Before treatment, REC sites had substantially lower accessibility and H3K27ac than RET and DIS sites (**Figure 2G-H**). DEX produced pronounced increases in both signals at REC sites (median paired changes +2.92 and +1.77 VST units), compared with much smaller changes at RET and DIS sites. Notably, H3K27ac at REC sites increased after DEX to approximately the level observed at untreated RET sites. Together, these data support a model in which GR-associated LEF1 recruitment marks GC-dependent activation of previously weak or inactive distal elements, whereas RET and DIS sites represent pre-existing active elements with more modest remodeling (**Figure 2I**).

### DNA sequence and chromatin accessibility differentially predict GR+LEF1 regulatory states

Because the three GR+LEF1 classes differ in enhancer state, we next asked whether their divergent responses were associated with differences in underlying DNA sequence. Motif analysis distinguished the GR+LEF1 classes (**Figure 3A-B**; **Table S3**). REC sites were strongly GRE-enriched: 80.5% contained a GR motif versus 6.5% of LEF1-only sites (adjusted odds ratio=50.8), and GR motifs were enriched relative to GR-only sites (adjusted odds ratio=24.5). In contrast, TCF/LEF and RUNX motifs increased from REC to DIS, and DIS sites were enriched for additional B-cell regulatory sequence features (PU.1/SPIB, EBF, and AP-1/ATF/CREB). RET sites showed comparatively few class-specific motif enrichments. Thus, REC sites show a strong GR sequence signature, whereas DIS sites retain features of pre-existing B-cell regulatory elements.

**Figure 3.**
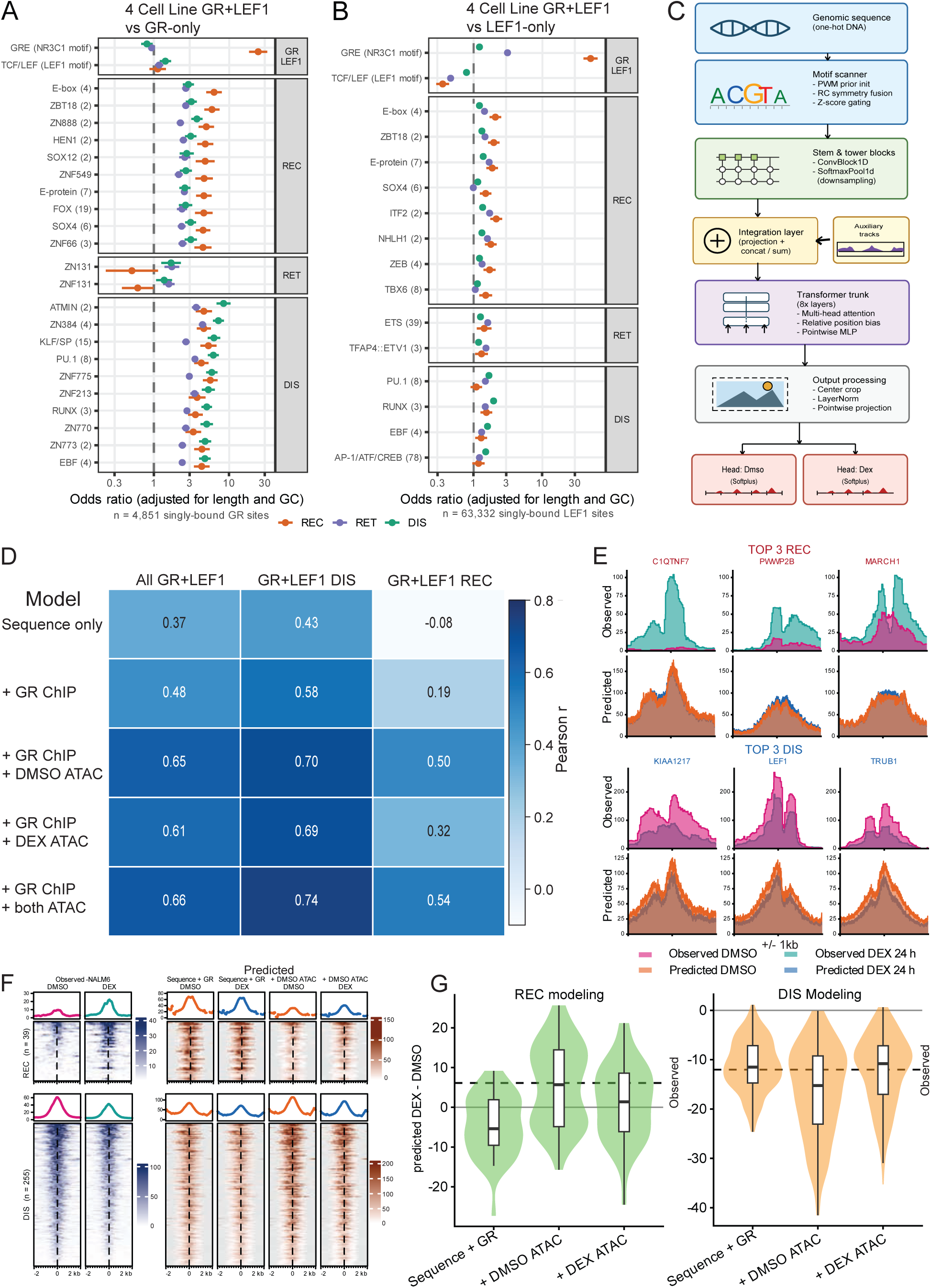
DNA sequence and chromatin accessibility differentially predict GR+LEF1 regulatory states. **(A-B)** Motif enrichment scores across GR+LEF1 recruitment (REC), retention (RET), and dissociation (DIS) sites compared with (A) GR-only or (B) LEF1-only sites. Motifs for transcription factors expressed in the analyzed B-ALL cell lines were evaluated using FIMO. Points indicate odds ratios adjusted for sequence length and GC content. GRE (NR3C1) and TCF/LEF (LEF1) motifs are shown together with the most enriched motif families for each GR+LEF1 class. **(C)** Schematic of the sequence-to-signal model used to predict GR+LEF1 regulatory states from genomic sequence with chromatin accessibility incorporated as an auxiliary input. Paired output heads predict DMSO- and DEX-associated signals. **(D)** Correlation heatmap summarizing model performance for prediction of LEF1 signal across all held-out GR+LEF1 sites and the DIS and REC subsets. Correlations are consistently lower for REC sites than for DIS sites, indicating that the recruitment response is less readily predicted by the evaluated sequence and auxiliary regulatory inputs. **(E)** Observed and predicted LEF1 CUT&RUN signal at the three REC and three DIS sites with the largest DEX-induced change on held-out chromosomes. Each panel spans one 2,048-bp model window; the nearest protein-coding gene is indicated (REC, red; DIS, blue). Predictions are from the sequence-plus-GR model. Observed and predicted signals are plotted on independent y-axes. **(F)** Observed and predicted LEF1 signal +/- 2 kb around REC (n = 39) and DIS (n = 255) sites on held-out chromosomes, ordered by observed DEX - DMSO change. Predictions are shown for the sequence-plus-GR model and for the same model with DMSO accessibility added. Observed and predicted signals are displayed on independent color scales, and the mean-signal plots above each heatmap are scaled independently for observed and predicted signal. Gray indicates positions outside the 2,048-bp model window. **(G)** Predicted DEX - DMSO change in LEF1 occupancy at REC sites (left; n = 39) and DIS sites (right; n = 255) for each model input: sequence-plus-GR, and the same model with DMSO or 24 h DEX accessibility added as an auxiliary track. Violins show site level predictions and boxes indicate the median and interquartile range. Dashed line, the observed median change at REC (+6.1) and DIS (-12.0).

We next asked whether broader sequence features could distinguish the GR+LEF1 response classes and whether incorporating chromatin accessibility improved prediction. We trained sequence-to-signal models to predict DMSO- and DEX-associated LEF1 occupancy from local sequence and GR signal, with chromatin accessibility incorporated as an auxiliary input in selected models (**Figure 3C**). Across held-out chromosomes, the models more readily predicted LEF1 loss at DIS than gain at REC sites (**Figure 3D-F**). Adding baseline or DEX-associated accessibility improved prediction, particularly at REC sites, but did not eliminate this difference (**Figure 3G**). These results indicate that dissociation is comparatively well associated with local sequence and pre-existing regulatory information, whereas recruitment is less readily predicted from sequence alone and is more dependent on chromatin context. This distinction is consistent with REC sites representing a regulatory transition in which GR engagement is followed by progressive chromatin opening and recruitment of additional factors.

### LEF1 recruitment is associated with progressive chromatin accessibility and coordinated regulatory-factor engagement

To define the chromatin dynamics accompanying LEF1 recruitment, we examined accessibility following DEX in NALM6 cells. Accessibility at REC sites increased progressively, with median log2 fold changes of +1.14, +1.70, and +2.81 at 1, 6, and 24 h, respectively (**Figure 4A-B**; **Figure S3A**). Accessibility increased at 92.5% of REC sites by 1 h and 97.9% by 6 h, indicating that subsequent increases largely reflect progressive strengthening at sites that responded early. In contrast, median accessibility changes at RET and DIS sites remained near zero over the early time course. Thus, chromatin opening at REC sites begins rapidly after GR activation and progressively increases following the early establishment of LEF1 recruitment.

**Figure 4.**
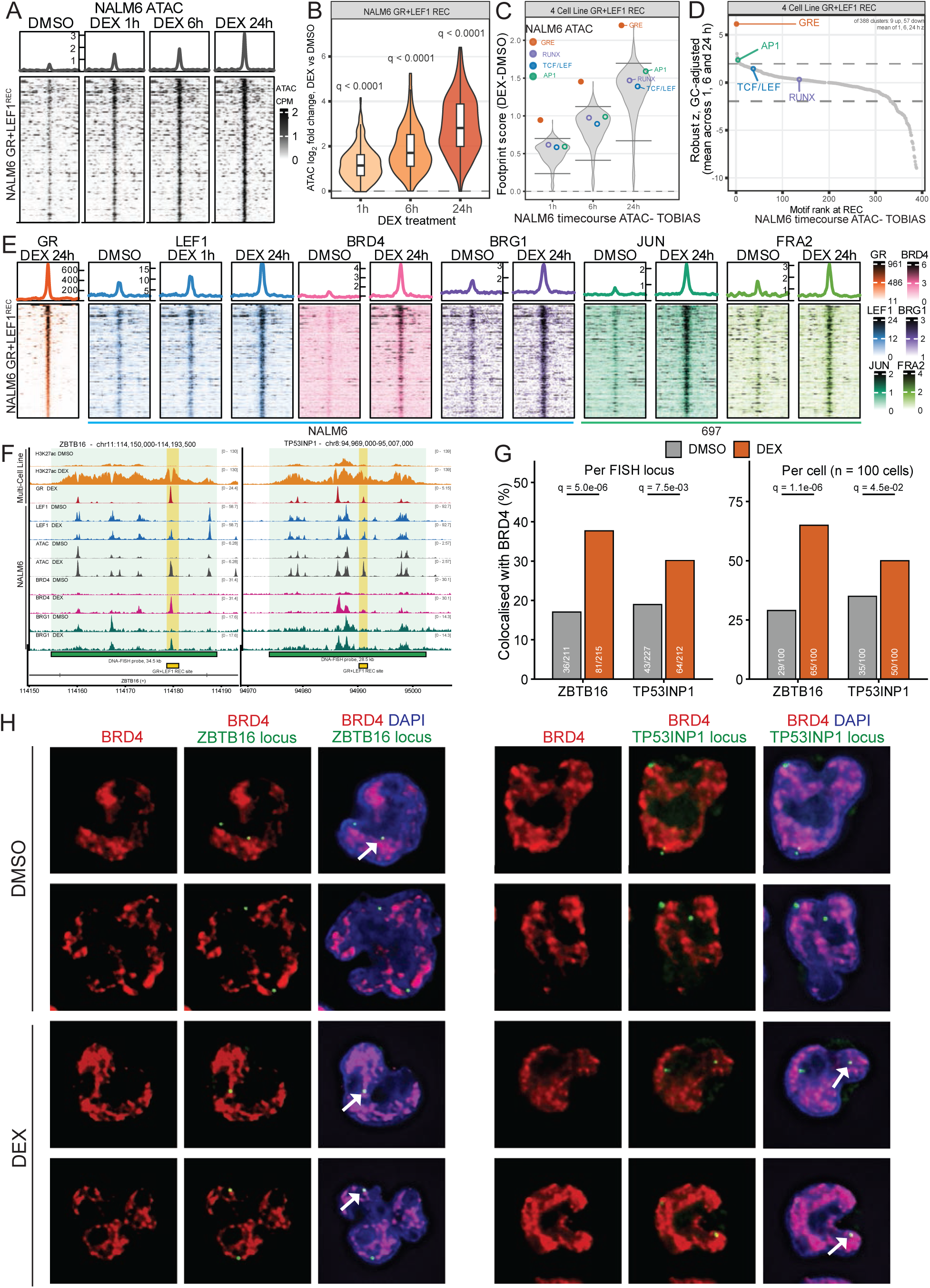
LEF1 recruitment is associated with progressive chromatin opening and coordinated regulatory-factor engagement. **(A)** ATAC-seq profiles and heatmaps at GR+LEF1 recruitment (REC) sites in NALM6 cells following 1, 6, and 24 h DEX treatment and matched DMSO controls. Rows represent the same REC sites across conditions. **(B)** DEX-induced changes in chromatin accessibility at REC sites across the ATAC-seq time course. Violin plots show log_2_ fold change relative to matched DMSO controls at 1, 6, and 24 h DEX. **(C)** TOBIAS differential footprinting at REC sites across the ATAC-seq time course. Violin plots show the distribution of DEX-induced differential footprint scores across all evaluated motifs. Points indicate representative GRE, TCF/LEF, RUNX1, and AP-1 motifs. **(D)** Ranking of DEX-induced differential footprint changes across 388 TOBIAS motif clusters at REC sites. Motif clusters with ≥100 instances across the 456 REC regions were included, resulting in 388 of 535 clusters meeting the inclusion threshold. Clusters were ranked by robust z scores summarizing differential footprinting across the three DEX time points. Dashed lines indicate z = ±1.96. The GRE cluster occupied the highest-ranked position. **(E)** Heatmaps and aggregate profiles at GR+LEF1 REC sites showing GR, NALM6 LEF1, BRD4 and BRG1 occupancy, and 697 JUN and FOSL2/FRA2 occupancy. BRD4, BRG1, JUN, and FOSL2/FRA2 are shown after 24 h DEX treatment and matched DMSO controls. Rows represent the same NALM6-defined REC coordinates and are ordered by mean BRD4 signal. **(F)** Genomic positions of DNA-FISH probes and GR+LEF1 REC sites at ZBTB16 locus and TP53INP1 region. **(G)** Spatial association of the ZBTB16 and TP53INP1 REC loci with BRD4-enriched nuclear domains following DMSO or 24 h DEX treatment. Left, percentage of DNA-FISH signals colocalizing with BRD4-enriched domains. Right, percentage of cells containing at least one colocalized FISH signal. Statistical comparisons of cell-level colocalization were performed using Fisher’s exact test with FDR correction where indicated. (H) Representative immunofluorescence/DNA-FISH images for the ZBTB16 locus (left) and TP53INP1 locus (right) following DMSO (top) or 24 h DEX (bottom) treatment. Arrows point to colocalized signals.

We next used ATAC-seq footprinting to examine inferred transcription-factor occupancy during this transition (**Figure 4C**; **Figure S3B-E**). At REC sites, the fraction of GR motif instances classified as bound increased from 12.5% in DMSO to 27.7%, 39.4%, and 54.6% after 1, 6, and 24 h DEX, respectively, whereas no comparable increase was observed at DIS sites. TCF/LEF, RUNX and AP-1 footprints also increased as REC sites became accessible. Thus, progressive chromatin opening at REC sites is accompanied by increased inferred occupancy of multiple TF families, with GR exhibiting a particularly strong response

We next ranked DEX-associated footprint changes of all motif clusters across the timecourse to determine which TF families showed the most robust and consistent enrichment or depletion over time. The GRE cluster ranked first at REC sites, whereas AP-1, TCF/LEF and RUNX motifs ranked 4^th^, 37^th^ and 136^th^, respectively (**Figure 4D**). GRE-family motifs also ranked highest at RET and DIS sites, whereas RUNX and AP-1 motifs ranked among the strongest negative footprinting responses at DIS sites (**Figure S3D-E**). This motif-ranking pattern was recapitulated across four B-ALL cell lines after 24 h DEX (**Figure S3F-G**), supporting a prominent GR-associated footprinting response together with distinct responses across additional TF families.

Because AP-1 footprints increased prominently as REC sites became accessible, we directly profiled JUN/cJUN and FOSL2/FRA2 occupancy by CUT&RUN after 24 h DEX treatment in 697 cells (**Figure 4E**). Both factors showed marked, significant gains at REC sites relative to vehicle control (JUN/cJUN: q=3.9e-48; FOSL2/FRA2: q=6.2e-36), providing direct occupancy evidence that the recruitment response extends beyond LEF1. Thus, GR-associated REC elements become sites of recruitment for multiple enhancer-associated transcription factors in addition to LEF1.

We next asked whether this broader regulatory-factor recruitment was accompanied by engagement of chromatin-remodeling and transcriptional coactivator machinery. We therefore examined recruitment of the chromatin remodeler BRG1 (SMARCA4) and the transcriptional coactivator BRD4 (**Figure 4E**; **Figure S3H**). Both factors increased significantly at REC sites following DEX treatment (BRD4: q=9.6e-46; BRG1: q=3.2e-40). Together with the LEF1 and AP-1 data, these findings indicate that REC sites undergo coordinated regulatory remodeling characterized by progressive chromatin accessibility and increased occupancy of multiple TFs, a chromatin remodeler, and a transcriptional coactivator.

Given the preferential increase in BRD4 occupancy at REC sites, we next asked whether this was accompanied by increased spatial association of REC elements with BRD4-enriched nuclear domains. We performed DNA FISH combined with BRD4 immunofluorescence at representative REC elements associated with *ZBTB16* and *TP53INP1*. Following 24 h DEX treatment, the fraction of *ZBTB16* REC-site FISH signals associated with BRD4-enriched nuclear domains increased from 17.1% to 37.7% (Fisher’s exact test, q=5.0e-6), while the *TP53INP1* REC site increased from 18.9% to 30.2% (Fisher’s exact test, q=7.5e-3; **Figure 4F-H**). The proportion of cells containing at least one colocalized signal also increased from 29% to 65% for the *ZBTB16* site (Fisher’s exact test, q=1.1e-6) and from 35% to 50% for the *TP53INP1* site (Fisher’s exact test, q=4.5e-2). Thus, orthogonal spatial analysis supports increased association of representative REC elements with BRD4-enriched nuclear domains following GR activation, further supporting coordinated regulatory remodeling at REC sites.

### RUNX1 dissociation parallels LEF1 redistribution following GR activation

The class-dependent RUNX footprinting pattern suggested that GR activation may reorganize RUNX-family transcription factor activity together with LEF1. Consistent with this observation, the RUNX1 motif was most frequent at DIS sites, occurring at 59.9% of DIS elements compared with 43.0% of RET and 37.9% of REC sites (**Figure 5A**). Differential footprinting across four B-ALL cell lines showed opposing RUNX-family responses at the GR+LEF1 site classes, with positive RUNX footprinting responses at REC sites and negative responses at DIS sites, the latter observed independently in each cell line (**Figure 5B-C**; **Figure S3E,G**; **Figure S4A**). Together, these findings implicated RUNX-family transcription factors in GC-induced regulatory reorganization and prompted us to directly examine RUNX1 occupancy.

**Figure 5.**
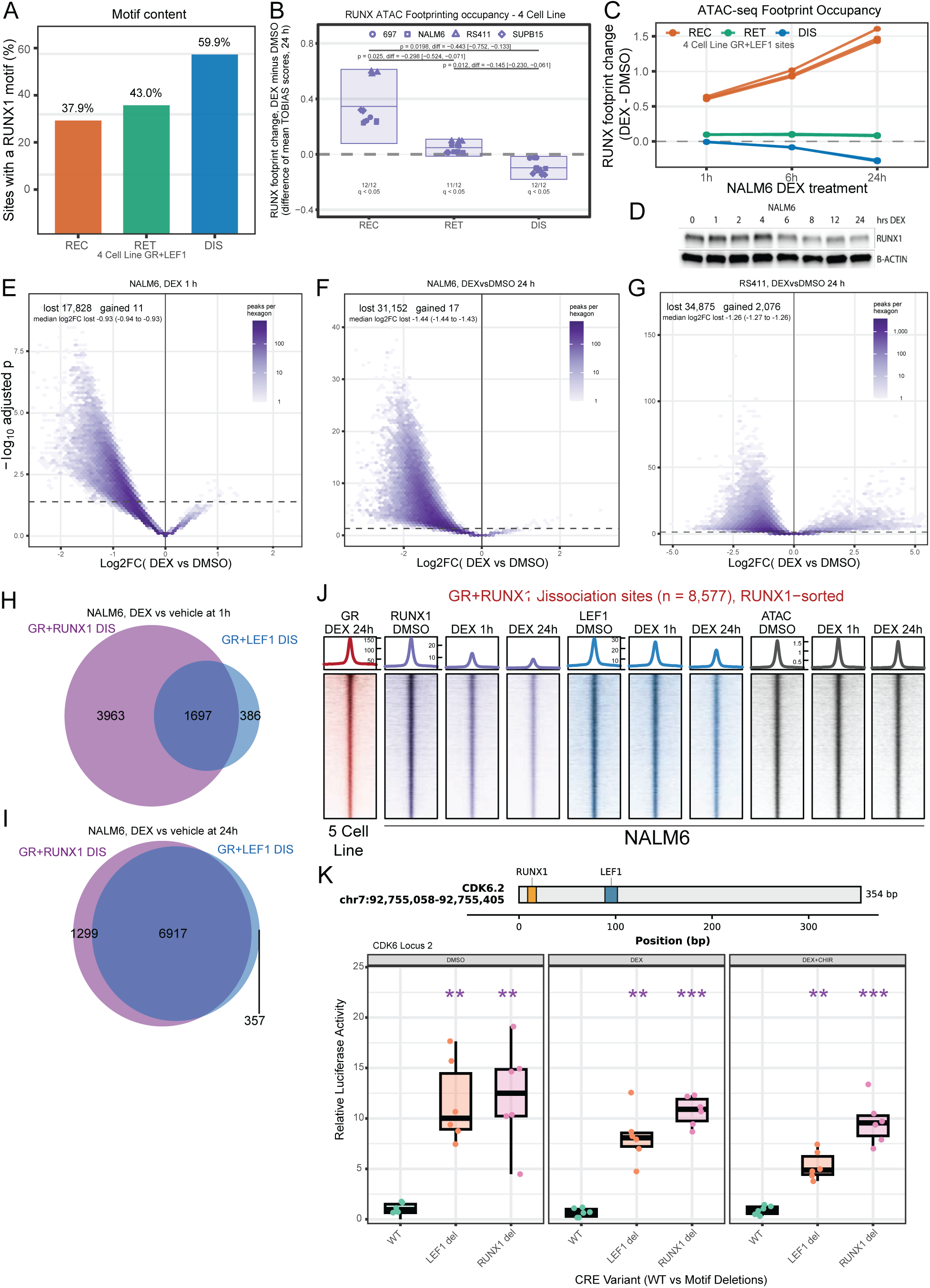
RUNX1 dissociation parallels LEF1 redistribution following GR activation. **(A)** Frequency of RUNX1 motifs at GR+LEF1 recruitment (REC), retention (RET), and dissociation (DIS) sites across four B-ALL cell lines. **(B)** Differential RUNX-family ATAC-seq footprint occupancy following 24 h DEX treatment across 697, NALM6, RS411, and SUPB15 cells. Points represent individual cell lines; boxes summarize the four-cell-line distribution. **(C)** RUNX-family footprint changes at REC, RET, and DIS sites across the NALM6 DEX time course relative to time-matched DMSO controls at the 4 cell line GR+LEF1 sites. **(D)** RUNX1 immunoblot following DEX treatment in NALM6 cells; β-actin serves as a loading control. **(E-G)** Differential RUNX1 CUT&RUN occupancy following 1 h **(E)** or 24 h **(F)** DEX treatment in NALM6 cells and 24 h DEX treatment in RS411 cells **(G)**. Numbers of significantly gained and lost sites and median log₂ fold changes are indicated. **(H-I)** Overlap between GR-associated RUNX1 and LEF1 dissociation sites in NALM6 cells following 1 h **(H)** and 24 h **(I)** DEX treatment. **(J)** Heatmaps of GR, RUNX1, LEF1, and ATAC-seq signal at GR+RUNX1 dissociation sites (n = 8,577), ordered by RUNX1 occupancy in vehicle-treated cells. **(K)** Luciferase activity of the CDK6.2 regulatory element containing wild-type (WT) sequence or targeted LEF1 and RUNX1 motif deletions following DMSO, DEX, or DEX+CHIR treatment. Reporter assays contained 5-6 replicate wells per construct and treatment. For panel B, filled points indicate significant within-cell-line differential footprinting after Benjamini-Hochberg correction (q < 0.05); brackets indicate paired comparisons across the four cell lines. For panel C, REC-versus-DIS comparisons were assessed by two-sided Mann-Whitney tests with Benjamini-Hochberg correction. For panel K, motif-deletion constructs were compared with the matched WT element and treatment effects were compared with DMSO-treated WT. FDR-adjusted significance is denoted *q < 0.05, **q < 0.01, and ***q < 0.001. Exact FDR values are shown where indicated.

We therefore profiled RUNX1 genomic occupancy and asked whether occupancy changes could be explained by altered protein abundance. RUNX1 protein remained stable through the first 4 h of DEX treatment (**Figure 5D**), whereas RUNX1 CUT&RUN revealed widespread loss of genomic occupancy. In NALM6 cells, 17,828 RUNX1 peaks were lost after 1 h DEX treatment compared with only 11 gained peaks, with the loss becoming more pronounced by 24 h (**Figure 5E-F**). A similarly predominant loss of RUNX1 occupancy was observed in RS411 cells at 24 h (**Figure 5G**). Thus, as observed for LEF1, changes in RUNX1 genomic occupancy precede detectable depletion of the protein.

RUNX1 and LEF1 dissociation occurred at substantially overlapping regulatory elements. GR-associated RUNX1 and LEF1 dissociation sites were enriched for overlap after 1 h DEX treatment (odds ratio=3.2, Jaccard index= 0.281, Fisher’s exact test, p=3.9e-92) and converged further by 24 h, when 6,917 sites were shared (odds ratio=5.4, Jaccard index=0.81, Fisher’s exact test, p=3.78e-91; **Figure 5H-I**). At GR-bound elements defined by RUNX1 loss, RUNX1 signal decreased by approximately 66% and LEF1 by 38%, from DMSO to 24 h DEX, whereas chromatin accessibility remained comparatively stable (**Figure 5J**). Together, these findings demonstrate that GR activation is accompanied by coordinated loss of RUNX1 and LEF1 from a shared set of pre-existing regulatory elements, rather than simply reflecting closure of the underlying chromatin.

To determine whether the sequence motifs occupied by these factors contribute to regulatory activity, we tested five GR- and LEF1-associated elements using luciferase reporters containing targeted motif deletions (**Figure 5K**; **Figure S4B-F**; **Table S4**). The elements exhibited distinct responses to DEX, providing diverse regulatory contexts in which to assess motif function (**Figure S4G**). RUNX motif deletion altered activity at each of the four elements in which RUNX motifs were tested, but the direction of the effect was element dependent. RUNX motif deletion reduced activity at CDK6.1 and STOML1, with combined RUNX motif deletion at STOML1 producing the strongest effect, whereas deletion increased activity at CDK6.2 and UCK2. GR motif deletion selectively impaired DEX-responsive activity at STOML1 and BLK, with comparatively little effect under vehicle conditions. TCF/LEF motif effects were similarly context dependent, with motif deletion increasing activity at both CDK6 elements and reducing activity under Wnt-activating conditions at UCK2, while effects at STOML1 and BLK were more modest. Thus, GR, TCF/LEF, and RUNX motifs make distinct, context-dependent contributions to regulatory element activity, supporting the functional importance of these sequence features within the regulatory elements undergoing GR-induced reorganization.

We next asked whether AP-1 occupancy also showed class-dependent behavior outside REC sites (**Figure S5**). In contrast to their prominent recruitment at REC sites, both JUN and FOSL2/FRA2 showed loss of occupancy at DIS sites following DEX treatment. DEX-reduced JUN peaks were significantly enriched at DIS sites (odds ratio=1.94, q=6.6e-20), whereas FOSL2/FRA2 showed broader genome-wide loss that included DIS sites. Thus, AP-1 exhibits opposing occupancy changes across REC and DIS regulatory classes, with recruitment at REC sites and dissociation from DIS sites. Together with redistribution of LEF1 and RUNX1, these findings support coordinated dissociation of multiple enhancer-associated TFs following GR activation.

### Wnt activation counter-regulates the glucocorticoid transcriptional response

Given the intersection between GR activation and the Wnt-associated regulatory network, we next asked how Wnt activation modifies an ongoing glucocorticoid response when both pathways are engaged simultaneously. RNA-seq following DMSO, DEX, CHIR, or DEX+CHIR treatment across four B-ALL cell lines revealed widespread and recurrent transcriptional antagonism (**Figure 6A-C**; **Figure S6A-C**). DEX altered 12,768 genes, whereas CHIR altered 4,812 genes and addition of CHIR to DEX altered 6,334 genes relative to DEX alone. Among 3,199 genes significantly regulated by both DEX and CHIR, combined treatment preferentially affected genes regulated in opposing directions (87.5% versus 25.5% in concordant quadrant; odds ratio=20.4, Fisher’s exact test, p=1.3e-261). This antagonistic response was observed across all four B-ALL cell lines, with 1,863 genes showing antagonism in at least three of four models.

**Figure 6.**
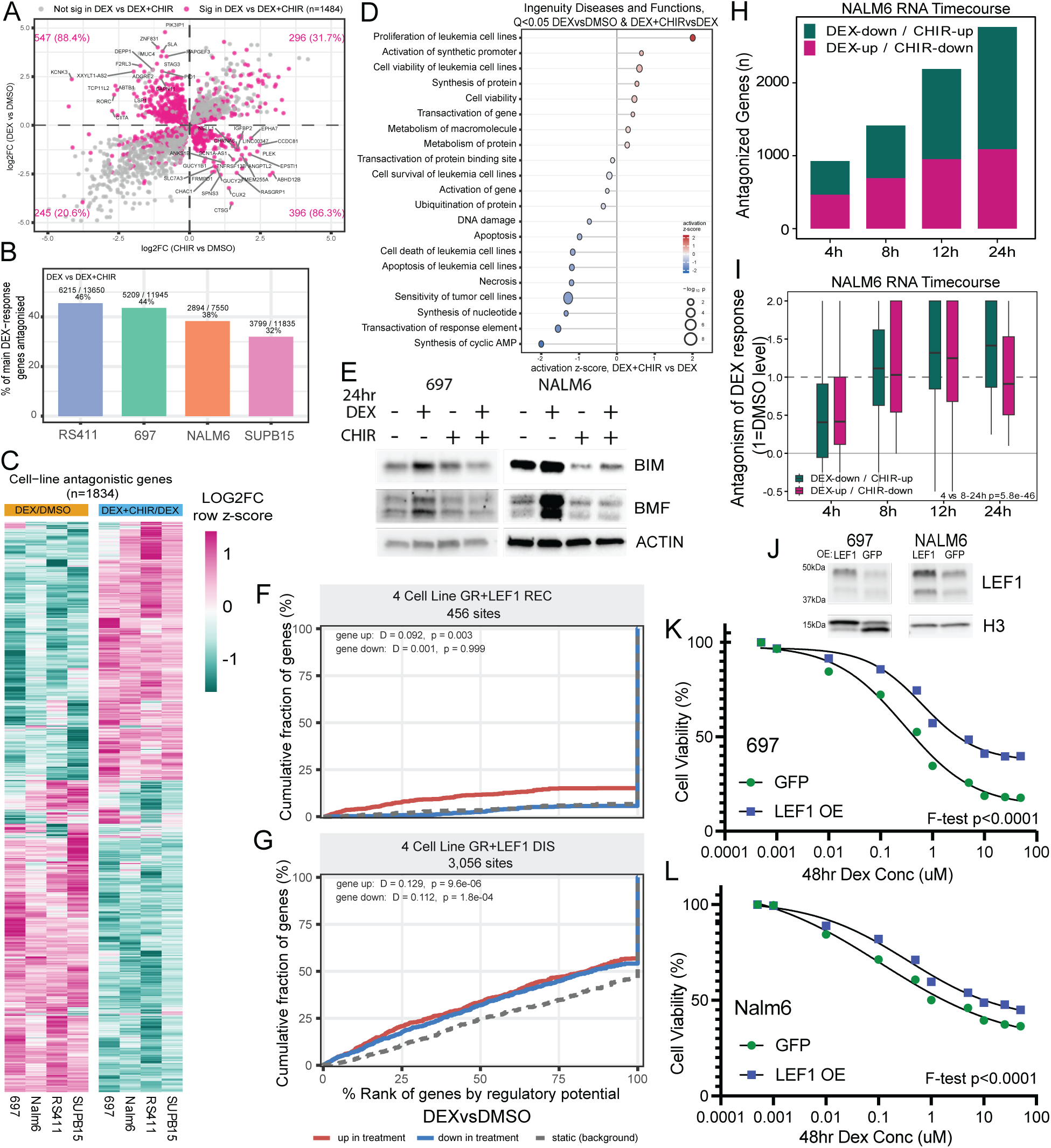
Wnt activation counter-regulates the glucocorticoid transcriptional response. **(A)** Comparison of log_2_ fold change gene-expression responses to 24 h dexamethasone (DEX) and CHIR99021 (CHIR) across 697, NALM6, RS411, and SUPB15 cells. The plot includes 3,199 genes significantly regulated by both DEX versus DMSO and CHIR versus DMSO; highlighted genes are additionally significantly altered by DEX+CHIR relative to DEX. Genes in opposing quadrants were significantly more likely to be altered by DEX+CHIR relative to DEX alone (88.4% of DEX up / CHIR down and 86.3% of DEX down / CHIR up) than genes in concordant quadrants (31.7% DEX/CHIR up, 20.6% DEX/CHIR down). **(B)** Frequency of transcriptional antagonism in individual B-ALL cell lines. Antagonism was observed in 45.5% of DEX-responsive genes in RS411, 43.6% in 697, 38.3% in NALM6, and 32.1% in SUPB15; 1,863 genes were antagonized in at least three of four cell lines. **(C)** Heatmap of recurrently antagonized genes across the four B-ALL cell lines. Row-standardized log₂ fold changes are shown for DEX versus DMSO and DEX+CHIR versus DEX. The 1,834 genes displayed were antagonized in at least three of four cell lines and had fold-change estimates in all eight comparisons. **(D)** Pathway enrichment results for recurrently antagonized genes are plotted according to enrichment and activation z-score. Ingenuity Disease and Functions annotations are provided. Proliferation of leukemia cell lines was predicted to increase (p = 9.0e-3, z = +2.01), whereas cell-death-related annotations were significantly enriched and consistently showed negative z-scores. **(E)** BIM and BMF protein abundance following 24 h treatment with DMSO, DEX, CHIR, or DEX+CHIR in 697 and NALM6 cells. β-actin serves as a loading control. **(F-G)** BETA analysis relating GR+LEF1 REC and DIS sites to DEX-responsive genes within the antagonized gene set. One-sided Kolmogorov–Smirnov tests were implemented by BETA. **(H)** Fraction of DEX-responsive genes antagonized by CHIR across a 4-24 h NALM6 time course. The antagonized fraction increased from 10.2% at 4 h to 14.6% at 8 h, 22.6% at 12 h, and 37.0% at 24 h. **(I)** Magnitude of CHIR-mediated antagonism during the NALM6 time course among 389 genes antagonized at three or more time points including 24 h. The fraction of the DEX response blocked by CHIR increased from a median of 0.416 at 4 h to 1.072, 1.280, and 1.107 at 8, 12, and 24 h, respectively. **(J)** Immunoblot confirming stable overexpression of full-length LEF1 relative to eGFP control in 697 and NALM6 cells. Histone H3 serves as a loading control. **(K-L)** Cell viability (y-axis) following 48 h DEX treatment across ten concentrations (x-axis) in cells stably expressing LEF1 or eGFP control. LEF1 overexpression increased survival in 697 **(K)** and NALM6 **(L)** cells.

Pathway enrichment analysis associated this antagonistic transcriptional program with increased leukemia-cell proliferation and reduced cell-death-related responses (**Figure 6D**). Among glucocorticoid-responsive apoptotic effectors, DEX induced the BH3-only genes *BCL2L11* (BIM) and *BMF*, whereas addition of CHIR attenuated their induction. BIM and BMF protein abundance showed corresponding changes, supporting antagonism of the glucocorticoid apoptotic program at both the transcript and protein levels (**Figure 6E**; **Figure S6D**). Consistent with this effect, Wnt activation reduced glucocorticoid sensitivity across all four B-ALL cell lines (**Figure S6E**).

We next asked whether GR+LEF1 sites undergoing LEF1 redistribution were associated with the antagonized transcriptional response. BETA analysis (34) linked both REC and DIS sites with DEX-induced genes within the antagonized gene set (**Figure 6F-G**). REC sites were significantly associated with DEX-induced but not DEX-repressed genes (p=2.9e-3), whereas DIS sites were associated with both DEX-induced (p=9.6e-6) and DEX-repressed (p=1.8e-4) genes. These findings connect GR-associated LEF1 redistribution with genes whose glucocorticoid response is opposed by Wnt activation.

To define the temporal development of this antagonism, we profiled NALM6 cells after 4, 8, 12, and 24 h of DEX, CHIR, and DEX+CHIR treatment (**Figure S6F-I**). The fraction of DEX-responsive genes antagonized by CHIR increased progressively from 10.2% at 4 h to 37.0% at 24 h (**Figure 6H**). Among 389 genes antagonized at multiple time points, CHIR blocked a median 42% of the DEX response at 4 h, whereas by 8 h the median response had returned approximately to the untreated level and remained strongly antagonized thereafter (paired t test, 4 versus 8-24 h, p=5.8e-46; **Figure 6I**). Importantly, GR protein loss under CHIR treatment was not apparent until after 12 h and was evident across all B-ALL cell lines at 24 h (**Figure S6J-K**). Thus, Wnt-mediated transcriptional antagonism begins before detectable GR protein loss and progressively strengthens over time.

Finally, because DEX reduces LEF1 abundance and Wnt activation opposes the glucocorticoid response, we directly tested whether increasing LEF1 abundance could alter glucocorticoid sensitivity. Stable LEF1 overexpression was confirmed in 697 and NALM6 cells (**Figure 6J**). LEF1 overexpression increased cell survival across the DEX dose response in both models, with the strongest effect observed in 697 cells (697, sum of squares F-test, p<1.0e-4; Nalm6, sum of squares F-test p<1.0e-4; **Figure 6K-L**). Together, these findings show that Wnt activation progressively counteracts the glucocorticoid transcriptional and apoptotic program during combined pathway activation, while LEF1 over-expression establishes that lineage-factor abundance is sufficient to modulate GC sensitivity.

### GR+LEF1 regulatory states are associated with glucocorticoid response and resistance in patient-derived B-ALL

We next asked whether the GR+LEF1 regulatory states identified in B-ALL cell lines were also evident in patient-derived leukemia. LEF1 CUT&RUN was performed in a primary B-ALL sample following 16 h DMSO or DEX treatment and evaluated across the GR+LEF1 recruitment (REC), retention (RET), and dissociation (DIS) sites defined in cell lines. Patient LEF1 occupancy was detected at 90.7% of DIS, 74.3% of RET, and 41.0% of REC sites (**Figure 7A-C**). Consistent with the cell-line-defined response classes, REC sites were enriched for DEX-specific LEF1 occupancy following DEX (**Figure 7A**). Quantitative analysis further demonstrated a pronounced decrease in LEF1 signal at DIS sites, an intermediate decrease at RET sites and a modest increase at REC sites following DEX (**Figure 7B-C**). Thus, the opposing behavior of REC and DIS sites identified in B-ALL cell lines was also evident in a primary B-ALL sample.

**Figure 7.**
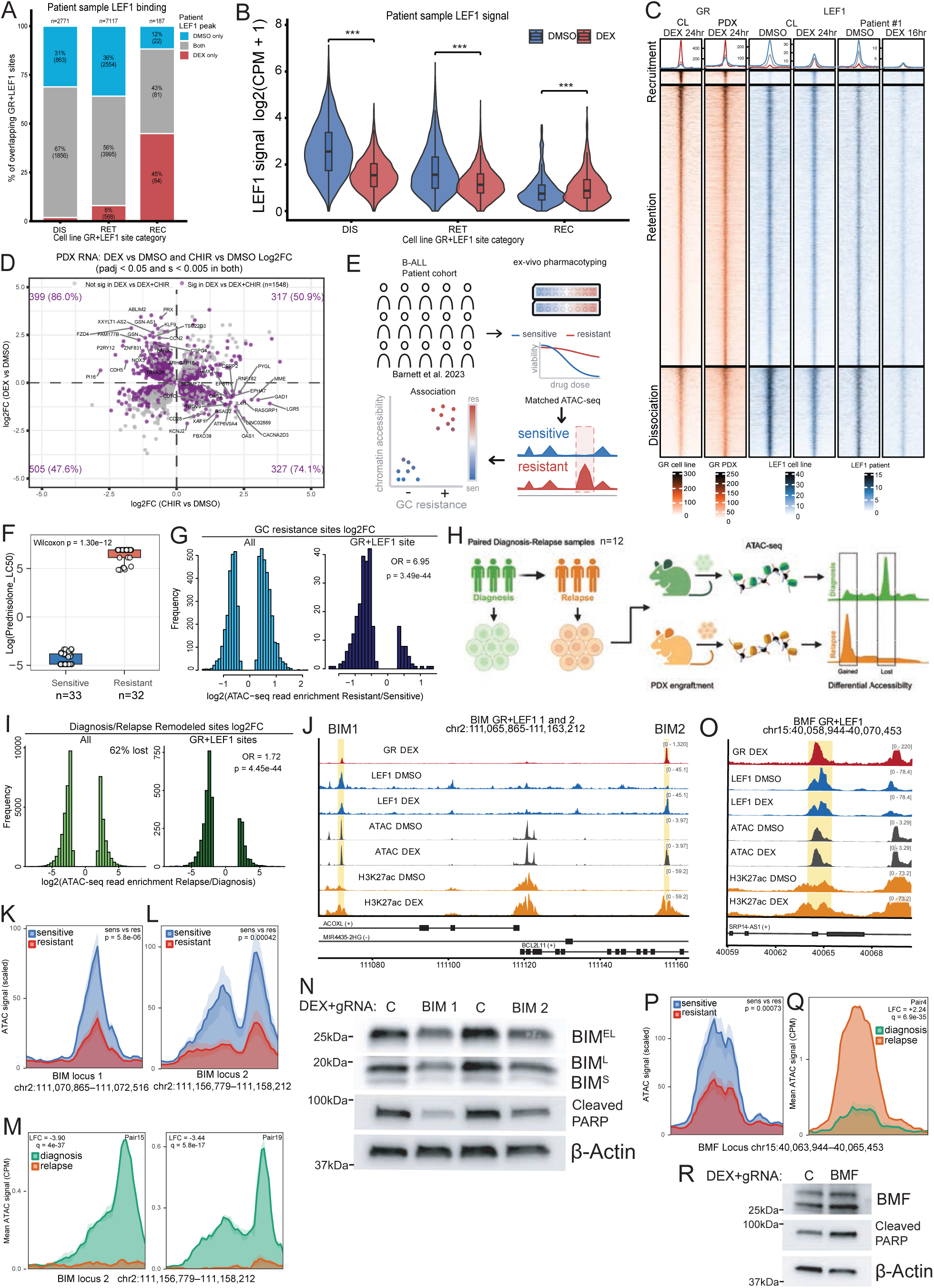
GR+LEF1 regulatory states are evident in patient-derived B-ALL and associated with glucocorticoid resistance and diagnosis-relapse remodeling. **(A)** LEF1 CUT&RUN peak composition at cell-line-defined GR+LEF1 dissociation (DIS), retention (RET), and recruitment (REC) sites in a primary B-ALL sample following 16 h DMSO or DEX treatment. Bars within each GR+LEF1 site class show the percentage of patient LEF1 sites identified in DMSO only, DEX only, or both conditions. Numbers indicate the number of patient LEF1 sites in at least one condition. **(B)** LEF1 CUT&RUN signal at DIS, RET, and REC sites in the same primary B-ALL sample following DMSO or DEX treatment. Signal is shown as log₂(CPM + 1). ***p < 0.0001, paired Wilcoxon signed-rank test with Benjamini-Hochberg correction. **(C)** Heatmaps of GR and LEF1 occupancy across GR+LEF1 REC, RET, and DIS sites. GR occupancy is shown in merged B-ALL cell-line and PDX datasets following DEX treatment; LEF1 occupancy is shown in B-ALL cell lines (left) and the primary B-ALL sample (right) following DMSO or DEX treatment. **(D)** Comparison of log_2_ fold change gene-expression responses to 24 h dexamethasone (DEX) and CHIR99021 (CHIR) across five B-ALL PDX models. The plot includes 2,589 genes significantly regulated by both DEX versus DMSO and CHIR versus DMSO; highlighted genes are additionally significantly altered by DEX+CHIR relative to DEX. Genes in opposing quadrants were significantly more likely to be altered by DEX+CHIR relative to DEX alone (86.0% of DEX up / CHIR down and 74.1% of DEX down / CHIR up) than genes in concordant quadrants (50.9% DEX/CHIR up, 47.6% DEX/CHIR down). **(E)** Schematic of the B-ALL patient cohort used to relate ex vivo prednisolone response to chromatin accessibility. Primary samples were pharmacotyped for prednisolone response, segregated by prednisolone LC50, and analyzed by matched ATAC-seq. **(F)** Prednisolone LC50 values for GC-sensitive (Q1; n = 33) and GC-resistant (Q4; n = 32) primary B-ALL samples. p = 1.30e-12, Wilcoxon rank-sum test. **(G)** Distribution of resistant-versus-sensitive ATAC-seq log₂ fold changes at GC resistance-associated sites genome-wide (left) and at the subset overlapping GR+LEF1 sites (right). Negative values indicate greater accessibility in GC-sensitive samples. **(H)** Schematic of the 12 matched diagnosis-relapse PDX pairs used to relate disease progression to chromatin accessibility. Diagnosis and relapse biospecimens were engrafted in mice as PDXs and harvested for ATAC-seq. Chromatin accessibility was compared within each diagnosis-relapse pair using DiffBind. Schematic was created with BioRender. **(I)** Distribution of the log₂ fold change (relapse versus diagnosis) among all strongly remodeled sites (| log₂FC|>2; left) and the subset overlapping GR+LEF1 sites (right). Negative values indicate reduced accessibility at relapse. Percentages indicate the proportion of sites showing accessibility loss at relapse. **(J)** Genome locus image of BIM locus 1 (BIM1) and BIM locus 2 (BIM2) GR+LEF1 sites. BIM1 is located upstream of *BCL2L11*/BIM and represents a GR+LEF1 RET site, whereas BIM2 is located within the *BCL2L11*/BIM gene locus and represents a GR+LEF1 REC site. **(K, L)** ATAC-seq coverage at two GR+LEF1 enhancers associated with BCL2L11 (BIM) in GC-sensitive and GC-resistant primary B-ALL samples: BIM locus 1 **(K)** and BIM locus 2 **(L)**. Lines represent mean accessibility and shading indicates SEM. p = 5.8e-6 and 4.15e-4, respectively, Wilcoxon rank-sum test. **(M)** ATAC-seq coverage at the BIM locus 2 enhancer in two matched diagnosis-relapse B-ALL pairs, both showing reduced accessibility at relapse: Pair 15 (log₂ fold change = −3.90; DiffBind q = 4.0e-37) and Pair 19 (−3.44; q = 5.8e-17). Log₂ fold changes are expressed as relapse relative to diagnosis. Lines represent mean accessibility and shading indicates SEM. **(N)** Immunoblot analysis of BIM and cleaved PARP in 697 cells following CRISPRi targeting of BIM locus 1 or locus 2 and treatment with DMSO or DEX. β-actin serves as a loading control. **(O)** Genome locus image of BMF GR+LEF1 RET site, which is located downstream of the BMF gene on the negative DNA strand. **(P)** ATAC-seq coverage at a GR+LEF1 regulatory element associated with BMF in GC-sensitive and GC-resistant primary B-ALL samples. Lines represent mean accessibility and shading indicates SEM. p = 7.25e-4, Wilcoxon rank-sum test. **(Q)** ATAC-seq coverage at the BMF regulatory element in a matched diagnosis-relapse B-ALL pair showing increased accessibility at relapse (log₂ fold change = 2.24; DiffBind q = 6.9e-35). Log₂ fold changes are expressed as relapse relative to diagnosis. Lines represent mean accessibility and shading indicates SEM. **(R)** Immunoblot analysis of BMF and cleaved PARP in 697 cells following CRISPRi targeting of the BMF regulatory element and treatment with DMSO or DEX. β-actin serves as a loading control.

We next determined whether the antagonistic relationship between GC and Wnt signaling was similarly observed in patient-derived models. RNA-seq across five B-ALL patient-derived xenografts following DEX, CHIR, or combined DEX+CHIR treatment revealed preferential antagonism of genes regulated in opposing directions by DEX and CHIR (**Figure 7D**), consistent with the B-ALL cell line response and also evident when cell-line and PDX RNA-seq datasets were analyzed together (**Figure S7A**). GC–Wnt antagonized genes were significantly enriched for overlap between cell lines and PDXs (odds ratio=3.62; Fisher’s exact test, p=4.5e-92), with 96.7% of shared genes exhibiting concordant directionality of the DEX response (**Figure S7B**). Thus, antagonism between GC and Wnt signaling is reproducible across B-ALL cell line and patient-derived models.

We then asked whether GR+LEF1 regulatory elements were associated with GC response in patients. Regulatory elements associated with GC resistance were identified using matched ATAC-seq and *ex vivo* prednisolone pharmacotyping from GC-sensitive (n=33) and GC-resistant (n=32) primary B-ALL samples (**Figure 7E-F**). Integration with GR+LEF1 elements identified 287 overlapped GC-resistance sites (**Table S5**). Whereas the 5,365 resistance-associated ATAC-seq sites showed a relatively balanced distribution of accessibility differences between GC-sensitive and GC-resistant samples, 84% of resistance-associated GR+LEF1 sites were more accessible in GC-sensitive samples, representing a significant enrichment compared with resistance-associated sites overall (odds ratio=6.95; Fisher’s exact test, p=3.49e-44; **Figure 7G**), a relationship observed across REC, RET, and DIS classes. These findings link the GR+LEF1 regulatory landscape to interpatient variation in GC response.

We next asked whether GR+LEF1 regulatory elements underwent substantial remodeling during disease progression using ATAC-seq from 12 matched diagnosis-relapse B-ALL pairs (**Figure 7H**; absolute log_2_ fold change >2; FDR<0.05; **Table S6**). Focusing on these robust accessibility changes, REC sites were preferentially remodeled between diagnosis and relapse (width-adjusted odds ratio=3.03; Wald test, BH-adjusted q=2.92e-25), whereas DIS sites were depleted for remodeling (width-adjusted odds ratio=0.65; Wald test, BH-adjusted q=5.02e-27).

Among strongly remodeled GR+LEF1 sites, all three classes showed a consistent bias toward loss of accessibility at relapse (**Figure 7I**, **Figure S7C**; odds ratio=1.72; Fisher’s exact test, p=4.45e-44). Notably, 142 of the 287 GC resistance-associated ATAC-seq sites overlapping GR+LEF1 elements (49.5%) were also strongly remodeled between diagnosis and relapse. These included elements associated with the pro-apoptotic GC effector genes *BCL2L11* (BIM) and *BMF,* linking GC response-associated regulatory elements to remodeling during disease progression.

Two GR+LEF1 enhancers associated with BIM showed significantly reduced accessibility in GC-resistant compared with GC-sensitive primary leukemias (**Figure 7J-L**). BIM locus 2 (REC site) showed markedly reduced accessibility at relapse in two matched patient pairs (**Figure 7M**), with significant but lower-amplitude decreases in two additional pairs (**Figure S7D**). Across the full cohort, accessibility at BIM locus 2 decreased in 9 of 12 pairs (Wilcoxon signed-rank test p=3.1e-2; **Figure S7E**). BIM locus 1 (RET site) did not meet the strong-remodeling criterion in any pair but showed significant, lower-amplitude decreases at relapse in three pairs (**Figure S7F**). CRISPRi targeting of either BIM enhancer reduced DEX-induced BIM protein and PARP cleavage, demonstrating that both elements contribute to the apoptotic GC response (**Figure 7N**; **Table S7**).

A GR+LEF1 element associated with BMF was likewise less accessible in GC-resistant primary leukemias (**Figure 7O-P**) and was identified among the GR+LEF1 elements remodeled during disease progression, with increased accessibility observed at relapse (**Figure 7Q**). Two additional diagnosis-relapse pairs showed significant but lower-amplitude decreases at relapse (**Figure S7G**), indicating that the direction of remodeling at this element varied across patients. CRISPRi targeting increased BMF expression and PARP cleavage following DEX, demonstrating that this resistance-associated element functions as a repressive regulator of BMF and modulates the apoptotic GC response (**Figure 7R**).

Together, these findings show that features of the GR+LEF1 regulatory landscape are evident in patient-derived B-ALL and are associated with interpatient variation in GC sensitivity and regulatory remodeling during disease progression. Functional perturbation of resistance-associated regulatory elements at BIM and BMF further demonstrates that components of this landscape regulate apoptotic effectors of the GC response.

## DISCUSSION

Our findings support a model in which canonical Wnt/LEF1 regulatory circuitry provides a central interface for GR-dependent reorganization of the B-ALL regulatory landscape. GR activation redistributed LEF1 into recruitment, retention, and dissociation states with distinct chromatin responses: dissociation preferentially involved pre-existing Wnt-responsive elements, whereas recruitment occurred at largely distinct GR-associated elements that acquired enhancer features. RUNX1 showed a parallel, predominantly dissociative response at many of the same pre-existing elements, while direct profiling of JUN and FOSL2/FRA2 demonstrated class-dependent AP-1 redistribution, including pronounced recruitment at LEF1-defined REC sites. The early LEF1 and RUNX1 occupancy changes began before detectable depletion of either protein, arguing that the initial transition is not a secondary consequence of reduced factor abundance. Together, our findings define how therapeutic GR activation reorganizes the endogenous B-ALL regulatory landscape into distinct transcription-factor occupancy and enhancer states that extend across multiple regulatory factors and are associated with therapeutic response.

REC sites illustrate one arm of this reorganization. Previous work established that a subset of steroid-receptor binding events is strongly encoded by receptor recognition sequences and can be shared across cellular contexts; Gertz et al. showed that shared steroid-receptor sites are distinguished by high-affinity response elements, whereas cell-type-specific sites depend more strongly on local chromatin and cooperating factors (35). Our data extend that framework in B-ALL by determining how this sequence-favored class of GR sites participates in reorganization of the leukemia lineage regulatory network. GR+LEF1 REC elements are strongly enriched for GREs, are comparatively weak or inactive before treatment, and become progressively accessible, with recruitment of LEF1, JUN, FOSL2/FRA2, BRD4, and BRG1. Representative REC elements also show increased spatial association with BRD4-enriched nuclear domains after DEX. Thus, these sites are not merely newly occupied GR elements; they can function as focal points at which receptor engagement is coupled to recruitment of multiple enhancer-associated factors and coactivator machinery, linking sequence-directed GR binding to broader remodeling of the leukemia lineage regulatory network.

DIS sites reveal a complementary arm of this process. These elements are enriched for TCF/LEF, RUNX, AP-1/ATF/CREB, and other B-cell regulatory sequence features and are strikingly enriched for β-catenin occupancy after Wnt activation, placing them within a pre-existing lineage-associated regulatory program. LEF1 and RUNX1 are coordinately lost from many of these elements even though chromatin accessibility remains comparatively stable at most DIS sites. Direct profiling of JUN and FOSL2/FRA2 further demonstrates loss of AP-1 occupancy at DIS sites following GR activation, extending this dissociative response across multiple enhancer-associated TFs. Thus, GR activation can reorganize the TF composition of pre-existing lineage-associated regulatory elements without requiring wholesale closure or decommissioning of the underlying chromatin. The context-dependent effects of RUNX, TCF/LEF, and GR motif disruption in reporter assays further indicate that the functional consequences of this redistribution reflect local regulatory grammar rather than a uniform activating or repressive role for any single factor.

The temporal separation between rapid LEF1/RUNX1 redistribution and later reductions in protein abundance also suggests that the GC response contains mechanistically distinct phases. GR binding can occur within minutes and is followed by sequential changes in cofactors, histone modifications, chromatin accessibility, and transcriptional output (17). In this framework, early redistribution of LEF1 and RUNX1 may represent an early chromatin-level response to GR activation, whereas later changes in lineage-factor abundance could reinforce or stabilize the remodeled state. Similarly, Wnt-associated transcriptional antagonism precedes later GR protein reduction, arguing that GR loss is unlikely to initiate the response but may contribute to its maintenance or amplification.

The regulatory intersection between GR and the Wnt-associated network was evident at both transcriptional and phenotypic levels. Across four B-ALL models, genes regulated in opposing directions by DEX and Wnt activation were strongly enriched for a response to combined treatment, and this antagonism accumulated over time. Wnt activation attenuated induction of pro-apoptotic effectors including BIM and BMF and reduced glucocorticoid sensitivity, whereas enforced LEF1 expression increased survival during DEX treatment. Clinical studies have likewise linked LEF1 expression to divergent outcomes in B-ALL, with high LEF1 associated with unfavorable outcome in adult cohorts but favorable prognosis in a pediatric cohort (28–31). These apparently divergent observations suggest that the consequences of LEF1 abundance may depend on developmental, molecular, or treatment context, consistent with our finding that LEF1 participates in distinct regulatory states following GR activation. The same transcriptional relationship was reproduced in B-ALL PDX models. These findings connect the lineage network reorganized by GR to the therapeutic glucocorticoid response while also indicating that GC-Wnt antagonism is not adequately explained by a single proximal mechanism.

Recent work showing tight β-catenin control and leukemia-cell death after pharmacologic GSK3β inhibition provides context for Wnt regulation in B-ALL (36). Our use of CHIR was primarily perturbational: it allowed us to identify Wnt-responsive regulatory elements and test how Wnt activation modifies the GC response. The observation that β-catenin stabilization can be detrimental to B-ALL does not diminish the regulatory antagonism observed here, but it does emphasize that the consequences of Wnt-pathway manipulation are context- and treatment-dependent. This distinction may be important when considering β-catenin-stabilizing strategies in combination with glucocorticoids.

The primary leukemia data further connect this regulatory architecture to therapeutic response. GR+LEF1 elements overlapping GC resistance-associated sites were strongly biased toward greater accessibility in GC-sensitive primary samples. A similar directional bias was observed among substantially remodeled GR+LEF1 elements during disease progression, which preferentially lost accessibility at relapse. The propensity for substantial remodeling differed across regulatory classes, with REC sites enriched and DIS sites depleted. Together, these findings link reduced accessibility at GR+LEF1 elements to both GC resistance and relapse while revealing class-specific differences in their susceptibility to substantial remodeling. One BIM element recovered here was previously characterized as a lymphocyte-specific GR/CTCF-regulated enhancer whose accessibility is reduced in GC-resistant ALL PDXs (37), and was subsequently implicated in PU.1-associated GC-dependent regulatory remodeling (19), providing an independent benchmark for the broader regulatory network identified here. We extend that locus-specific observation by placing BIM within a genome-wide class of GR-associated elements undergoing lineage-factor redistribution, linking related elements to GC response across a larger primary B-ALL cohort, identifying relapse-associated remodeling, and functionally interrogating BIM and BMF elements. The convergence of regulatory redistribution, primary patient chromatin states, disease progression, and functional perturbation supports a model in which components of the redistributed network contribute to GC-induced apoptosis.

Several limitations should be considered. ATAC-seq footprinting provides motif-centered inference rather than direct TF occupancy, although CUT&RUN provides orthogonal support for RUNX1, JUN, and FOSL2/FRA2 redistribution. While rapid LEF1 recruitment to GR-bound, GRE-rich elements supports a GR-centered mechanism, the molecular intermediates coupling GR activation to redistribution of LEF1, RUNX1, AP-1, and cofactors remain to be defined. RET sites may also be heterogeneous, encompassing both stable elements and cell-type-specific changes that do not meet the aggregate differential threshold. LEF1 overexpression establishes that LEF1 abundance can modulate GC sensitivity but not that every component of the GR+LEF1 network is required for therapeutic response, and CHIR has biological effects beyond canonical Wnt activation. Finally, although the patient and relapse analyses connect this regulatory architecture to clinically relevant states, additional longitudinal and perturbational studies will be needed to establish the contribution of individual elements to acquired resistance in vivo.

In summary, our study establishes that the response of B-ALL cells to therapeutic GR activation is shaped by dynamic reorganization of the endogenous regulatory landscape rather than engagement of a fixed set of regulatory elements. By integrating transcription-factor redistribution with chromatin, cofactor, transcriptional, functional, and patient derived data, we connect these regulatory state transitions to GC response and resistance. The convergence of GC-Wnt transcriptional antagonism, altered regulatory states in primary leukemia, and remodeling during disease progression further supports the therapeutic relevance of this architecture. Together, these findings provide a framework for understanding how steroid-receptor signaling interacts with endogenous leukemia regulatory circuitry to shape drug response.

## METHODS

### Cell Culture

Human B-ALL cell lines NALM6 (DSMZ ACC128), 697 (DSMZ ACC42), RS411 (DSMZ ACC508), SUPB15 (DSMZ ACC389) and SEM (DSMZ ACC546) cells were used for experimentation. All cell lines were routinely tested for mycoplasma contamination and maintained in RPMI 1640 (ThermoFisher, 21870084) supplemented with 10% FBS and 1% penicillin/streptomycin/GlutaMAX (ThermoFisher, A5873601). Cultures were maintained at 37 °C and 5.0% CO₂ in a humidity-controlled incubator. CUT&RUN, ATAC-seq, ChIP-seq, and RNA-seq experiments were performed by treating cell lines at a density of 1.5e6 cells per ml with appropriate compounds. Unless otherwise indicated, cells were treated with 1 µM dexamethasone (Cell Signaling Technologies, 14776), 3 µM CHIR99021 (Cell Signaling Technology, 54290) or both compounds. DMSO was used as vehicle control.

### Patient biospecimens

Primary patient samples were obtained from St. Jude Children’s Research Hospital, and patients or their legal guardians provided the required written informed consent. The use of patient biospecimens was approved by the institutional review board at St. Jude Children’s Research Hospital.

### Western Blotting

Cells were washed with ice-cold PBS, then lysed in RIPA lysis buffer (ThermoFisher, 89901) supplemented with a protease inhibitor cocktail (Sigma, P8340). Five to 15 µg of whole-cell extract was resolved on 4–15% Tris-Glycine gradient gels (BioRad 4561085), transferred onto nitrocellulose membranes (BioRad, 1620213), blocked in 5% milk in TBST (Tris-Buffered Saline with 0.1% Tween-20), and incubated with primary antibodies overnight at 4 °C. Primary antibodies were used against GR (BD Biosciences, 611227), LEF1 (Cell Signaling Technology, 2230), *β*-Catenin (Cell Signaling Technology, 8480), RUNX1 (Abcam, ab240639), BCL2L11 (Cell Signaling Technology, 2819), BMF (Cell Signaling Technology, 50542), cleaved PARP (Cell Signaling Technology, 5625), β-actin (Cell Signaling Technology, 4970; loading control) and Histone H3 (Cell Signaling Technology, 9715; loading control). Membranes were washed with TBST, incubated with secondary antibody, and then imaged using the BioRad ChemiDoc and analyzed with BioRad ImageLab.

### Cell line drug response assays

Cell viability following treatment with dexamethasone and CHIR99021 was measured using CellTiter-Glo^®^ 2.0 Cell Viability Assay (Promega, G9243) according to the manufacturer’s protocol. Cells were plated on 96-well plates at 2e4 cells per well and treated with varying concentrations of drug for 24, 48 and/or 72 h. Luminescence was measured on a BioTek Cytation1 (Agilent). Experiments were performed using 3 biological replicates and at least 3 technical replicates per drug concentration. Statistical significance was measured by sum of squares F-test comparing drug response curves against the null hypothesis that the data could be explained by 1 curve. Curves were fitted and compared in GraphPad Prism 11; p values below 0.0001 are reported as p<1.0e-4.

### CUT&RUN experimentation

CUT&RUN experiments were performed using the CUTANA ChIC/CUT&RUN Kit (EpiCypher, 14-1048, v6.0) with 5e5 cells per reaction according to the manufacturer’s protocol. Antibodies and quantities per reaction were as follows: 2 µL (1:25, 12 ng) of β-Catenin (D10A8) Rabbit Monoclonal Antibody (Cell Signaling Technology, 8480), 1 µL (0.5 µg) of Anti-RUNX1 / AML1 antibody [EPR23044-100] (Abcam, ab240639), 1 µL (1:50, 0.157 µg) LEF1 (C12A5) Rabbit Monoclonal Antibody (Cell Signaling Technology, 2230), 1 µL (1:50, 34.5 ng) of FRA2 (D2F1E) Rabbit Monoclonal Antibody (Cell Signaling Technology, 19967), 2.5 µL (0.5 µg) of JUN/c-Jun antibody (EpiCypher, 13-2019), 2.5 µL (0.5 µg) of BRG1 antibody (EpiCypher, 13-2002), 0.5 µL (0.5 µg) of BRD4 antibody (EpiCypher, 13-2003). β-Catenin – 4 cell lines per treatment, FRA2 – 2 replicates DMSO vs 3 replicates DEX), JUN/c-Jun - 3 replicates per treatment, RUNX1-RS411, NALM6 – 4 replicates per timepoint/treatment, LEF1 timecourse NALM6 – 4 replicates per timepoint/treatment, LEF1 4 cell line – NALM6, 697, SUPB15 3 replicates, RS411 2 replicates per treatment, LEF1 – patient sample, 1 replicate per treatment. replicates were collected and processed from individual wells for each condition. CUT&RUN libraries were sequenced on an Illumina NovaSeq X Plus by the Hartwell Center at St. Jude Children’s Research Hospital.

### ATAC-seq

A total of 1e5 cells per replicate and 4 replicates per condition were processed using the ATAC-seq kit (Active Motif, 53156) according to the manufacturer’s protocol. ATAC-seq libraries were sequenced on an Illumina NovaSeq X Plus by the Hartwell Center at St. Jude Children’s Research Hospital. ATAC-seq data from primary patient leukemia specimens was obtained from our previously published study (38).

### ChIP-seq

ChIP-seq was performed as previously described (38) on 2-2.5e7 cells with minor modifications described here. Formaldehyde-crosslinked cells were lysed in Cell Lysis Buffer III for ChIP (bioWORLD, 10450055) supplemented with 1x protease inhibitor tablet (Roche, 11836170001). Chromatin samples were incubated with antibodies for GR (BD Biosciences, 611227) or H3K27ac (Abcam, ab4729) as follows. For each sample, 200 µL of magnetic IgG beads (GR, Invitrogen, Dynabeads™ M-280 Sheep Anti-Mouse IgG, 11202D; H3K27ac, Invitrogen, Dynabeads™ M-280 Sheep Anti-Rabbit IgG, 11204D) were washed three times in binding buffer (1x PBS + 5 mg/ml BSA [fraction V] + 1x protease inhibitor cocktail) and resuspended in 1 ml binding buffer. Five micrograms of anti-H3K27Ac antibody (Abcam, ab4729) was added to each magnetic bead solution, which was rotated overnight at 4°C. H3K27Ac was performed on 697, NALM6, RS411, and SUPB15 as biological replicates. GR was performed on 697, NALM6, RS411, and SUPB15 and SEM and 6 PDX samples as biological replicates. ChIP-seq libraries were sequenced on an Illumina NovaSeq X Plus by the Hartwell Center at St. Jude Children’s Research Hospital.

### RNA-seq

Total RNA was isolated from 2-3 replicates per cell sample using the RNeasy Plus Mini Kit (Qiagen, 74134) from approximately 2-3e6 cells. RNA was processed using the Illumina stranded total RNA prep kit. RNA-seq libraries were sequenced on an Illumina NovaSeq X Plus by the Hartwell Center at St. Jude Children’s Research Hospital.

### LEF1 over-expression

Lentiviral *LEF1* (GenScript Clone ID: OHU23027; Accession No.: NM_016269.5) and control e*GFP* overexpression constructs cloned into a pLVV-EF1a-P2A-Puro plasmid backbone were obtained from GenScript. The Vector Laboratory at St. Jude Children’s Research Hospital generated lentiviral particles of LEF1 and eGFP over-expression plasmids. Cells were plated in 12-well plates at 2e6 cells per well in complete RPMI supplemented with polybrene (8 µg/mL) and transduced into NALM6 or 697 cells. Cells were washed 24 h after transduction and puromycin selection (NALM6 = 1 µg/mL; 697 = 2 µg/mL) began 72 h post-transduction and continued for 7 days prior to drug viability assays using Cell-Titer Glo.

### Luciferase reporter assays

Plasmid constructs (see **Table S4**) were inserted into pGL4.23 (Promega, E841A) upstream of the minimal promoter. Cloned pGL4.23 and pRL-TK Renilla plasmid constructs were co-transfected into 697 cells using the Neon transfection system (Thermo Fisher Scientific, #MPK5000, 10μL Neon tips; 1600V, 10ms, 3 pulses; 1μg plasmid DNA and 0.1μg pRL-TK control vector). For each biological replicate, 2.5e5 cells were transfected and allowed to incubate for 24 h post-transfection. After 24 h, transfected cells were treated for 24 h with DMSO vehicle control, 1 µM dexamethasone (Cell Signaling Technologies, 14776) or 3 µM CHIR99021 (Cell Signaling Technology, 54290). Luciferase activity was measured in 96-well plates using the Dual Luciferase Reporter Assay System (Promega, E1960) on a BioTek Cytation1 (Agilent). A total of 5-6 replicate wells from independent transfections were utilized per construct. Statistical significance was measured by ANOVA with multiple testing comparisons.

### ImmunoFISH

ImmunoFISH experimentation was performed by the CytoStem Shared Resource at St. Jude Children’s Research Hospital. NALM6 samples were treated with DMSO or DEX for 24 h for immunofluorescence (IF) for BRD4 & DNA FISH with two test site probes (ZBTB16, and TP53INP1) to assess the spatial relationship between BRD4-enriched nuclear domains and the test probe loci. Anti-BRD4 Alexa Fluor 647 conjugated rabbit monoclonal antibody (Abcam, ab197608) was used as the antibody at a dilution of 1:100. For the DNA FISH assays purified ZBTB16 fosmid DNA (WI2-908N21 / 11q23.2 / chr11:114,149,083-114,190,497 hg38), and TP53INP1 fosmid DNA (WI2-2759L24 / 8q22.1 / chr8:94,963,946-95,005,241 hg38) were labeled separately with a green-dUTP (Seebright 496) by nick translation. Single cells from each sample were independently applied to glass slides by cytocentrifugation. Slides were fixed in 1% PFA in PBS for 5 minutes followed by 1% PFA in PBS plus 0.05% Igepal for 5 minutes, and stored in 70% ETOH at -20°C. The slides were rinsed in PBS and blocked in 200µl blocking buffer (1% BSA, 2X SSC) and then incubated with the Anti-BRD4 antibody at 1:100 dilution in antibody diluent for 45 minutes at RT in the dark. Slides were washed in PBS and stained with Vectashield containing DAPI and imaged for IF. Slides were then washed in PBS and digested with RNaseA for 45 minutes at 37°C. Cross linking with 4% PFA was then performed, followed by treatment in 0.2N HCL. Slides were denatured in 70% formamide, 2X SSC at 80°C for 8 minutes followed by ETOH dehydration series. Denatured DNA probes were then added in a solution containing sheared human cot DNA with 50% formamide, 2X SSC, 10% dextran and hybridized overnight at 37°C. Slides were then washed in 50% formamide, 2X SSC at 37°C for 5 minutes and stained with Vectashield containing DAP1 and imaged again using the same coordinates as had been imaged for the IF images. Imaging was performed in 3D using a widefield fluorescence microscope (Nikon Eclipse Ti2) equipped with Nikon Nis Elements version 6.02 software and Autoquant 3D deconvolution. Images were deconvolved using the Richardson-Lucy with number of rounds set to auto-select. Images were acquired using 0.1-micron (0.1 µm) plane spacing and a 60X planapochromatic objective with a numerical aperture of 1.4. Captured images were analyzed using in-house Z-Fisher software, a custom image-analysis platform developed and maintained by the CytoStem Shared Resource. The software was used to identify and quantify immunofluorescence and DNA-associated fluorescent signals and to determine the spatial relationship between signal centroids. Colocalization was defined based on the three-dimensional distance between the immunofluorescence and DNA signal centroids, with a maximum distance threshold of 0.65 µm. For each analyzed nucleus, the software calculated the frequency of DNA signals exhibiting immunofluorescence colocalization relative to the total number of DNA signals detected. This standardized, automated analysis was applied across experimental conditions to minimize operator-dependent variability and provide consistent quantitative measurements of immunofluorescence-DNA spatial association.

### Pharmacotyping

The sensitivity of primary leukemia cells to prednisolone was determined ex vivo for a total of 65 pediatric or adult patients with ALL. Patients were enrolled on a treatment protocol at St. Jude Children’s Research Hospital (TOTXVI; NCT00549848), (TINI; NCT02553460) or on ECOG-ACRIN or CALGB protocols. Patients and/or their guardians provided written informed consent in accordance with the Declaration of Helsinki. The study was approved by the Institutional Review Board (IRB) of St. Jude Children’s Research Hospital.

Ex vivo prednisolone sensitivity was determined in primary ALL cells using a four-day MTT viability assay, as previously described (5). Following drug exposure, MTT was added to a final concentration of 0.45 mg/mL, and cultures were incubated for an additional 6 h. Formazan generated by viable cells was solubilized in acidified isopropanol and measured spectrophotometrically. Prednisolone LC50 values were estimated using a four-parameter dose-response model.

### CRISPR interference screening

697 cells expressing dCas9-KRAB were generated by transduction with the lentiviral vector lenti-dCas9-KRAB-blast (Addgene, 89567) at an MOI of 10. Transduced cells were maintained in 1 µg/mL blasticidin S (Research Products International, B12150) for at least 7 days. dCas9-KRAB-expressing cells were then transduced at an MOI of 10 with a lentiGuide-Puro (Addgene, 52963)–derived pool of three guideRNAs per locus. GuideRNA sequences were designed with GuideScan2 (https://guidescan.com/py/grna_design) using a specificity cutoff of 0.2. gRNA-transduced cells were selected for at least 7 days with 0.5 µg/mL puromycin (GenDEPOT, CR026) and 1 µg/mL blasticidin S before assays. To assay the effect of CRISPRi on drug response, 1e6 cells/mL dCas9-KRAB cells (control) and dCas9-KRAB-gRNA cells were cultured with 1 µM dexamethasone (Cell Signaling Technologies, 14776) or vehicle control (DMSO) for 24 h prior to western blotting.

### Machine Learning

Training examples were defined as 2,048-bp genomics windows centered on peak summits (+/- 1024bp). Peaks separated by less than 1,024 bp were deduplicated. Windows located outside primary chromosomes or overlapping the ENCODE hg38 blacklist were excluded. The data were split by chromosome, with two chromosomes reserved for validation, three for testing, and the remaining chromosomes for training. DNA sequences were one-hot encoded, auxiliary bigWig tracks were extracted at base-pair resolution, and target signals were summed into non-overlapping 2-bp bins, yielding 1,024 bins per window.

**Model architecture** The model follows the general architectural design of Enformer (39), comprising a convolutional encoder, a transformer trunk and multiple linear output heads. It further incorporates an interpretable motif-scanning input layer and an optional pathway for integrating auxiliary covariates measured at base-pair resolution.
**Motif-scanning input layer** In the motif-based configuration, the one-hot sequence is passed through a fixed convolutional motif scanner initialized using Position weight matrices (PWMs) from a MEME-format file. PWM values were converted to log-odds scores against a uniform nucleotide background, mean-centered, and standardized to a 29-bp kernel width. Reverse-complement filters were added to produce two times number of motifs fixed filters, whose weight remained frozen during training. Scanner outputs were rectified and summed across forward and reverse-complement orientations to yield strand-invariant motif activations. Channels whose maximum score did not exceed a background-derived z-score threshold of 3.0 were suppressed. The resulting motif activation map replaces the one-hot sequence as model input. Alternatively, the raw one-hot sequence can be passed directly to the convolutional stem.
**Convolutional encoder layer** The input is projected to 256 channels by a width-15 convolution, followed by a residual block with group normalization, width-5 convolutions, GELU activations, and dropout. Attention pooling performs downsampling, while channel width is doubled after every second block. The number of blocks is automatically determined by the target output resolution; for a 2,048-bp input and 1,024-bin output, a single block provides two-fold downsampling.
**Auxiliary covariate integration** When auxiliary per-base tracks are supplied, they are linearly interpolated to the downsampled length of the convolutional representation, projected to the same channel width by a 1×1 convolution, and combined with the sequence representation either by concatenation along the channel axis (default) or by addition.
**Transformer and output layers** The representation is projected to 512 dimensions and processed by eight transformer blocks with 8-head self-attention, relative positional biases, and 2,048-dimensional feed-forward layers. After center-cropping to 1,024 bins, a pointwise transformation is applied, followed by task-specific linear heads with softplus activation to produce non-negative predictions.
**Model training and evaluation** The model was trained with a variance-normalized, masked mean squared error loss. Optimization was performed with AdamW with a learning rate of 1e-5 and weight decay of 1e-2. Gradients were clipped to a maximum norm of 1.0. Depending on the dataset, the models were trained for 20-30 batch epochs with batch size ranging from 4 to 8. The checkpoint with the lowest validation loss was retained for test-set evaluation. The best validation checkpoint was evaluated on held-out test chromosomes. Performance was assessed for each output head using mean squared error (MSE) and Pearson correlation across valid bins. Training and evaluation were performed on Cray XD670 nodes, each with eight NVIDIA H100 SXM GPUs (80 GB VRAM per GPU), 112 CPU cores and 1 TB RAM; each job used a single H100 GPU.
**Held-out chromosome analyses** The held-out chromosome analyses were restricted to the test chromosomes not used for model fitting or validation. REC and DIS sites on these chromosomes were evaluated separately to compare model performance across regulatory classes. Representative loci were selected by the magnitude of the observed DEX-associated LEF1 change, and aggregate observed-versus-predicted profiles were summarized across held-out REC and DIS sites. Predictions from models incorporating accessibility were compared with the corresponding sequence-plus-GR configuration to assess the contribution of chromatin context.

### Identification of Prednisolone LC50 associated accessible chromatin

ATAC-seq data from B-ALL patient samples were processed with nf-core ATAC-seq pipeline version 2.1.2. Genomic regions of interest were defined using merged ATAC-seq signal across the patient cohort as input for MACS2 peak calling (40). These merged and filtered MACS2 peak regions were used as input to Subread for ATAC-seq read counts matrix generation across the ATAC-seq patient cohort. In preparation for linear modeling prednisolone LC50 values for each patient sample were log transformed, z-scaled and filtered for only the upper and lower quartile cohorts (upper quartile = prednisolone resistant, lower quartile = prednisolone sensitive). Differential chromatin accessibility associated with prednisolone LC50 was identified using DESeq2 modeling ATAC-seq read counts against categorical classification of prednisolone LC50 values with the Wald statistical test. Significant differential chromatin accessibility was defined as p-adjusted < 0.05. Log fold change effect sizes were shrunk using apeglm to deflate log fold change values for low read count regions.

### Patient-derived xenografts and Diagnosis-Relapse Analysis

All animal studies were approved by the Institutional Animal Care and Use Committee of St. Jude Children’s Research Hospital. Patient-derived xenografts (PDXs) were expanded as previously outlined (41). Briefly,12 diagnosis-relapse pairs (#4, 7, 8, 9, 11, 12, 14, 15, 17, 18, 19, 20) from the Children’s Hospital of Philadelphia were expanded in female NSG mice (NOD.Prkdc^scid^Il2rd^tm1Wjl^/SzJ, 8-12 weeks old). PDX cells (1-2e6) were injected intravenously via the tail vein and animal health was monitored regularly. Engraftment was monitored by peripheral-blood flow cytometry and cells were stained with mTER119 (BioLegend, 116228; diluted at 1:200), mCD45 (BD Biosciences, 557659; 1:100), hCD45 (BD Biosciences, 555482; 1:50) and hCD19 (BioLegend, 363004; 1:100), and the percentage of hCD45 and hCD19 positive cells were evaluated using a BD FACS LSR II machine (BD FACS Diva Software v.9). PDX cells were harvested from spleen when leukemia cells reached 80% in peripheral blood or when mouse became moribund.

ATAC-seq was performed as described, with two technical replicates per PDX sample. Differential ATAC-seq sites were identified using DiffBind (42) within each of the 12 diagnosis-relapse pairs using absolute log_2_ fold-change >2 and FDR<0.05. Sites meeting these criteria across pairs were merged to generate a nonredundant set of remodeled regions. All ATAC-seq sites across pairs were similarly merged to generate a background universe used for enrichment analysis.

### STARR-seq analysis

STARR-seq data were obtained from our previous publication, with GC-responsive STARR-seq elements identified as previously described (22).

### Processing of NGS data

All initial processing of NGS data was performed using nextflow nf-core pipelines: atacseq 2.1.2, chipseq 2.1.0, cutandrun 3.2.2, using the HG38 genome.

### Peak calling

Peaks for GR, LEF1, β-catenin, RUNX1, JUN and FOSL2 were called with Genrich at q < 0.05 on query-name-sorted, replicate-merged alignments, run in replicate mode separately for each factor and treatment group. Summits were taken from the Genrich narrowPeak offset, using the strongest overlapping peak per interval. Interval arithmetic used bedtools v2.31.0 and v2.31.1 (43).

### Differential occupancy and chromatin accessibility

Fragment counts over consensus peaks were obtained with featureCounts (subread v2.1.1) in paired mode with -O and --fracOverlap 0.2, and tested in DESeq2 (44) in R v4.4.0 with a negative binomial generalized linear model and a Wald test on the treatment contrast. Fold changes were shrunk with apeglm where s-values are reported. A peak is called changed at adjusted p < 0.05 together with an s-value < 0.005, and the direction is taken from the sign of the shrunken fold change. This rule is applied identically to every differential binding and accessibility analysis in the paper. Peak-based size factors divide out any response that approaches genome-wide, which applies to many factors after DEX. The hg38 genome was tiled into 100 kb windows with bedtools makewindows, every called peak was padded by 1 kb and combined with the blacklist, any window intersecting that union was discarded, and 20,000 peak-free bins were sampled. Size factors were estimated on the background count matrix with DESeq2::estimateSizeFactorsForMatrix and assigned to the peak DESeqDataSet directly; peak-based estimation was not called. The same factors, inverted, are the deepTools scale factors used for the corresponding coverage tracks, so tracks and statistics share one scale. All CUT&RUN data were normalized the same way. CUT&RUN libraries were assessed on read depth, on the fraction of read pairs in peaks and on the median inter-replicate Spearman correlation. The NALM6 accessibility time course was fitted as a single DESeq2 model over all replicate libraries on one 160,469-peak Genrich consensus.

### Per-site occupancy change at defined sites

Where a factor is summarized at a fixed set of sites rather than at its own peaks, background-scaled coverage was averaged over the central window at each site and the DEX minus DMSO difference tested by two-sided Wilcoxon signed rank, with the Hodges-Lehmann shift and a 95% confidence interval as the effect size. A pseudocount is added per factor, taken as the 10th percentile of that factor’s own non-zero coverage, because a single pooled value shrinks fold changes on low-background tracks harder than on high-background tracks.

### Differential gene expression analysis

All three RNA-seq datasets, the four-cell-line panel, the NALM6 time course and the xenograft panel, were analyzed under one rule. Counts were fitted in DESeq2, fold changes were shrunk, and a gene is called changed at adjusted p < 0.05 together with an s-value < 0.005, with no fold-change threshold. The same rule was applied to DEX versus DMSO, CHIR versus DMSO and DEX+CHIR versus DEX.

### Motif analysis

Motifs were scanned with FIMO (MEME Suite v5.5.4 and v5.5.7) in --text mode at a single fixed threshold of p < 10⁻⁴ applied identically to every sequence set, against a database of 997 motifs from JASPAR2026 and HOCOMOCO H14CORE restricted to factors expressed in these cell lines (TPM > 0.2). The sequence sets differ in median length and in median GC fraction, and both inflate motif detection independently of biology. Enrichment was therefore fitted as a logistic model of motif presence on class, log length and GC fraction. Effect sizes are the exponentiated class coefficient with a Wald 95% confidence interval, and p values are Benjamini-Hochberg corrected within each control. Singly bound sites were split against the merged GR and LEF1 peak sets into LEF1-only (63,332), GR-only (4,851). Each class was tested separately against the LEF1-only and the GR-only control, because a pooled control is 89.4% LEF1-only and is in effect a LEF1-only control. The two contrasts ask what co-binding adds relative to a LEF1 site and relative to a GR site. Motifs were grouped by matrix similarity rather than by name, because name-based grouping is unreliable. TOMTOM was run all against all, similarity taken as the capped negative log p, symmetrized, converted to a distance and clustered by average linkage with the tree cut at 0.80; clusters sharing a factor name were then merged. Across all full 997 motifs, this yielded 421 clusters and 360 families; among the 784 resolvable motifs used for reporting, this yielded 336 clusters and 286 families.

### Transcription Factor Footprinting

Footprinting used TOBIAS (45). ATAC-seq signal was corrected for Tn5 insertion bias with ATACorrect, footprint scores were computed with ScoreBigwig, and differential footprinting between conditions was computed with BINDetect over the scanned motif set, run separately within each site class so that scores are compared against motifs measured on the same regions. Bound fractions are the proportion of motif instances in a class that BINDetect classifies as bound in a given condition. The differential footprint rankings in Figure 4D and Supplemental Figure S3D-E were computed using the 535 motif clusters defined by TOBIAS BINDetect, which groups the 997 motif matrices independently of the TOMTOM-based grouping described above. These motif groupings were therefore treated as distinct analyses.

### Signal visualization and coverage heatmaps

Coverage matrices were computed with deepTools v3.5.6 computeMatrix reference-point, centered on the site, at ±2 kb or ±5 kb as stated in each legend, in 10 bp or 25 bp bins, with missing data treated as zero. Heatmaps were drawn with EnrichedHeatmap and ComplexHeatmap in R. Rows are sorted once, usually on vehicle signal of the factor named in the legend, and held in that order across every column of a panel.

### BETA analysis

Regulatory potential was computed with BETA against hg38 using the GR+LEF1 class BED files as the peak input, a distance limit of 250 kb and a peak limit of 15,000 and the DEX vs DMSO 4 cell line gene expression data.

### Statistics and reproducibility

All statistical tests were two-sided unless stated otherwise. The exception is the BETA analysis in Figure 6F-G, where the Kolmogorov-Smirnov statistic implemented by BETA is one-sided by construction. p values were corrected for multiple testing by the Benjamini-Hochberg procedure within each analysis, and corrected values are reported as q values or as adjusted p values. Exact p values are reported when available.

Central tendency and variation are reported as follows. Violin and box panels show the median and interquartile range. Coverage profiles show the mean across samples, with shading giving the standard error of the mean. For unadjusted enrichment analyses, odds ratios and 95% confidence intervals were estimated using Fisher’s exact test. Width-adjusted odds ratios and Wald 95% confidence intervals were obtained from logistic regression with site class and log10 peak width as predictors. Where significance was assessed with Pearson’s chi-square test, the reported odds ratios and confidence intervals were estimated separately using Fisher’s exact test. Per-site occupancy changes report the Hodges-Lehmann shift with a 95% confidence interval. Correlations report the Spearman rank correlation coefficient. The exact n for each comparison is given in the corresponding figure legend.

## ACKNOWLEDGEMENTS

We would like to thank the St.Jude Hartwell Center for next-generation sequencing of ATAC-seq as well as library preparation and next-generation sequence of CUT&RUN, ChIP-seq, and RNA-seq experimentation. We would also like to thank the Pharmacotyping Resource at St. Jude for performing *ex vivo* drug sensitivity assays on patient biospecimens as well as the St. Jude CytoStem shared resource for ImmunoFISH experimentation. This work was supported by the National Cancer Institute (R01CA234490, P30CA021765), the Human Genome Research Institute (HG013981), the National Institute of General Medical Sciences (P50GM115279) and the American Lebanese Syrian Associated Charities (ALSAC). The CytoStem shared resource is supported by the National Institute of Health, National Cancer Institute, Cancer Center support grant P30CA021765-46 (Cytogenetics) and ALSAC. The content is solely the responsibility of the authors and does not necessarily represent the official views of the National Institutes of Health.

## AUTHOR CONTRIBUTIONS

R.J.M. and D.S. conceived the study. R.J.M., K.R.Ba., K.R.C., S.B. and Y.C. developed the methodology. R.J.M., D.O., G.B., B.B., F.U., K.R.Bh., V.V., M.V., S.Y. and D.T.T. performed the investigation. R.J.M., D.O., G.B., M.Z., B.B., F.U., K.R.Bh., K.R.Ba., V.V., M.V., S.S., S.Y. and Y.C. performed the formal analysis. M.Z. and Y.C. developed the software. R.J.M. and K.R.Ba. curated the data and performed the validation. R.J.M., M.Z., K.R.Ba. and Y.C. prepared the visualizations. D.T.T. provided resources. R.J.M. and D.S. prepared the original draft. D.O., G.B., B.B., K.R.Ba., V.V., K.R.C., S.B. and D.S. reviewed and edited the manuscript. Y.C., J.J.Y. and D.S. supervised the study. All authors approved the final manuscript.

## MATERIALS & CORRESPONDENCE

Correspondence and requests for materials may be addressed to Daniel Savic.

## SUPPLEMENTAL FIGURE LEGENDS

**Figure S1.**
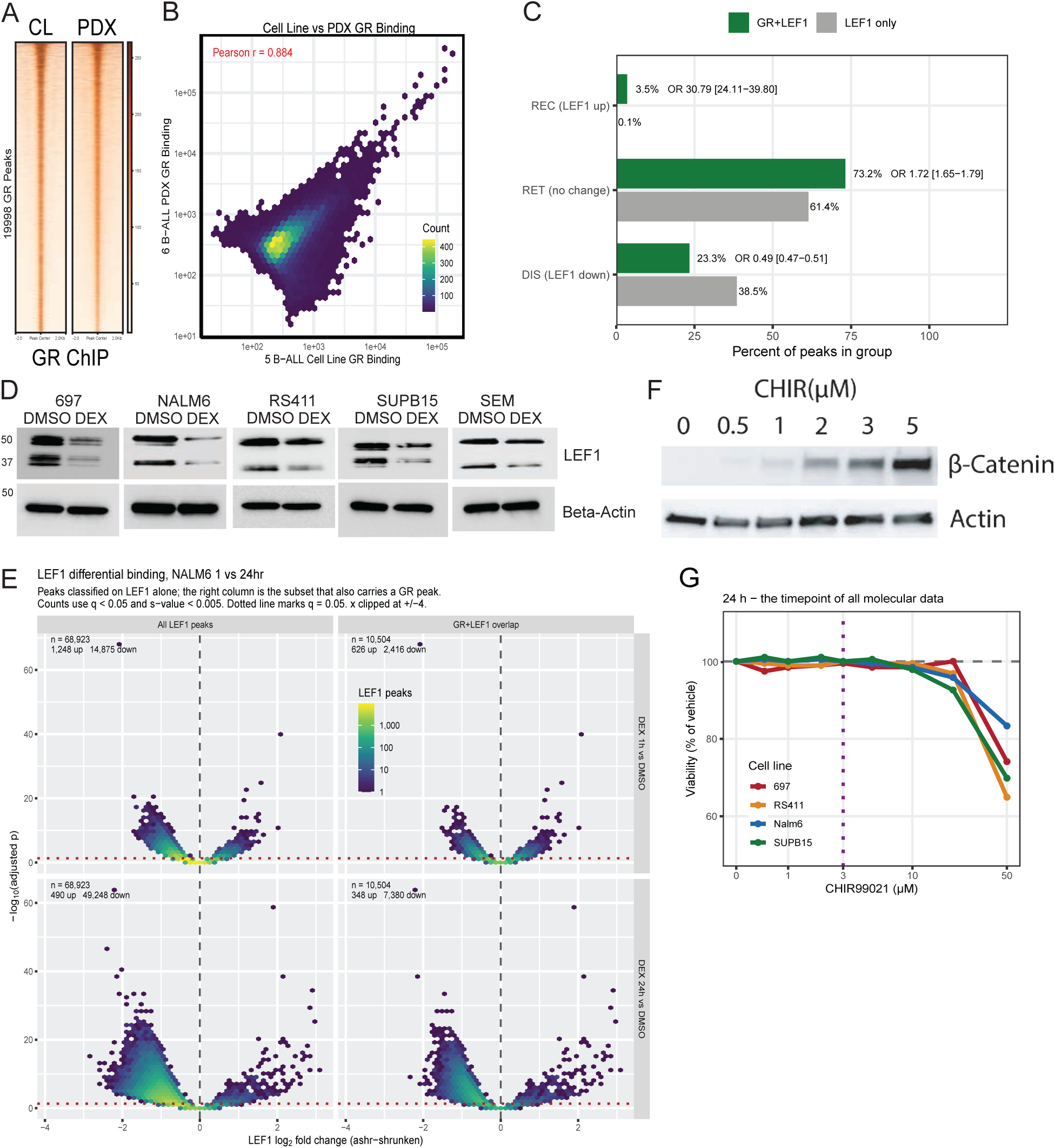
Characterization and controls supporting glucocorticoid-induced LEF1 redistribution. **(A-B)** Comparison of GR occupancy in cell lines (CL) and patient-derived xenografts (PDXs). Heatmaps of GR occupancy in both cell models is shown in **(A**). Correlation of GR occupancy between cell lines and PDXs is shown in **(B)**. **(C)** Comparison of LEF1 response to DEX at GR+LEF1 versus LEF1-only sites. Bars show the fraction of sites undergoing recruitment (REC), retention (RET), or dissociation (DIS). GR co-occupancy is strongly associated with LEF1 recruitment (odds ratio [OR] 30.79, 95% CI 24.11–39.80) and with reduced odds of LEF1 dissociation (OR 0.49, 95% CI 0.47–0.51). **(D)** Western blot of LEF1 protein abundance following 24 h treatment with vehicle (DMSO) or 1 µM dexamethasone (DEX) in 697, NALM6, RS411, SUPB15, and SEM cells. DEX reduces LEF1 protein across all five B-ALL cell lines. β-actin is shown as a loading control. **(E)** Volcano plots of differential LEF1 occupancy in NALM6 cells following 1 h (top) or 24 h (bottom) DEX treatment relative to DMSO. Analyses are shown for all 68,923 LEF1 peaks (left) and the subset of 10,504 LEF1 peaks overlapping GR (right). At 1 h, 1,248 sites gained and 14,875 lost LEF1 across all LEF1 peaks, whereas 626 gained and 2,416 lost LEF1 among GR-overlapping sites. At 24 h, 490 sites gained and 49,248 lost LEF1 across all peaks, whereas 348 gained and 7,380 lost LEF1 among GR-overlapping sites (NALM6-specific recruitment and dissociation sites shown in Figure 1F-G). **(F)** Western blot of β-catenin following treatment with increasing concentrations of CHIR99021, demonstrating dose-dependent stabilization of β-catenin. β-actin serves as a loading control. **(G)** Cell viability following 24 h treatment with increasing concentrations of CHIR99021 in 697, NALM6, RS411, and SUPB15 cells. The 3 μM concentration used for molecular profiling is indicated by the vertical dashed line.

**Figure S2.**
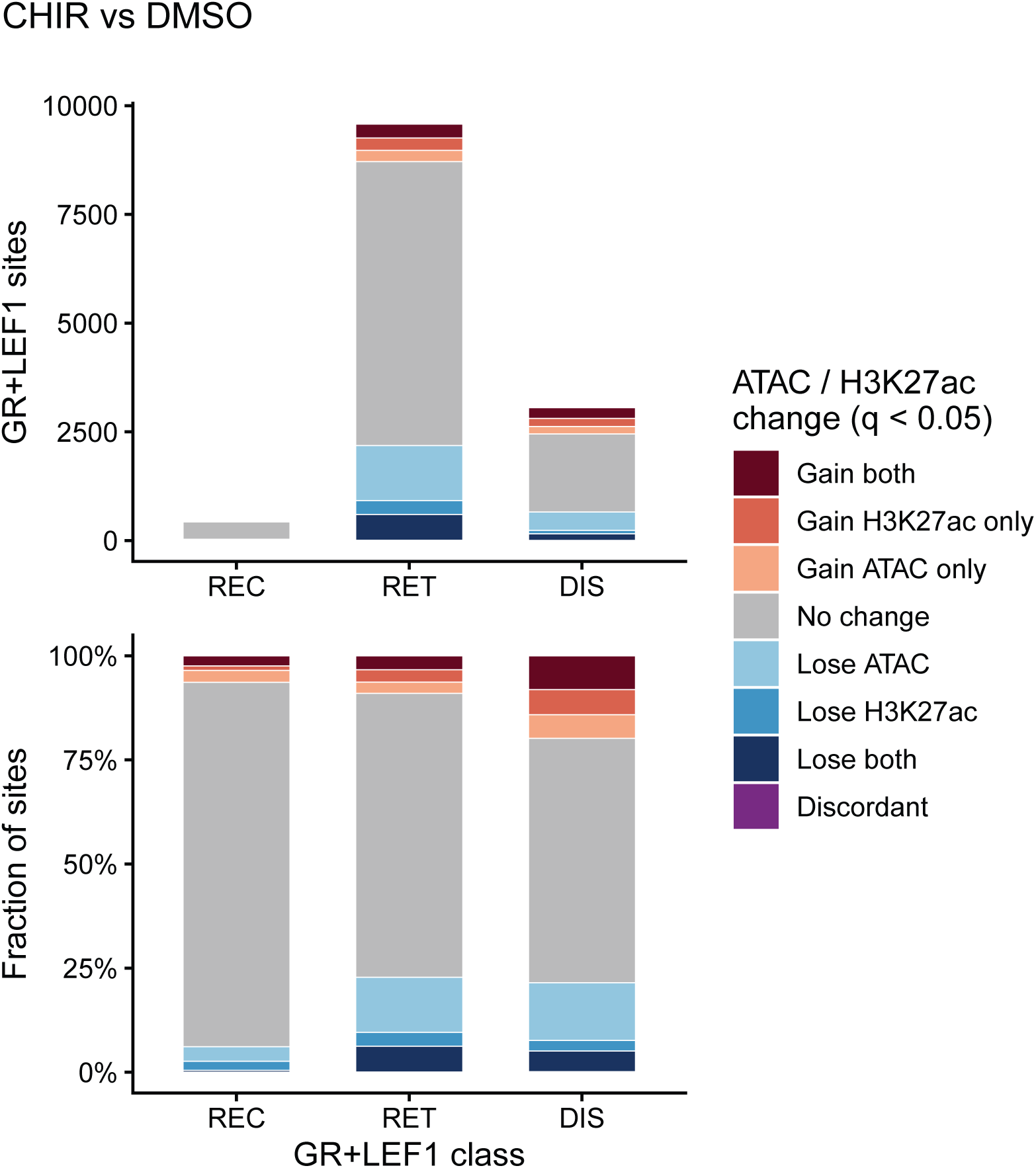
Wnt activation does not recapitulate the class-specific chromatin remodeling induced by glucocorticoids. Number (top) and proportion (bottom) of REC, RET, and DIS sites exhibiting significant changes in chromatin accessibility and/or H3K27ac following CHIR treatment relative to DMSO (q < 0.05). Sites were classified using the same criteria as in Figure 2B. Most REC sites remained unchanged following CHIR treatment (87.5%), with only 2.4% gaining both accessibility and H3K27ac, compared with 83.8% following DEX treatment. RET and DIS sites exhibited mixed gains and losses of chromatin accessibility and H3K27ac, demonstrating that Wnt activation does not reproduce the coordinated, class-specific chromatin response observed following DEX.

**Figure S3.**
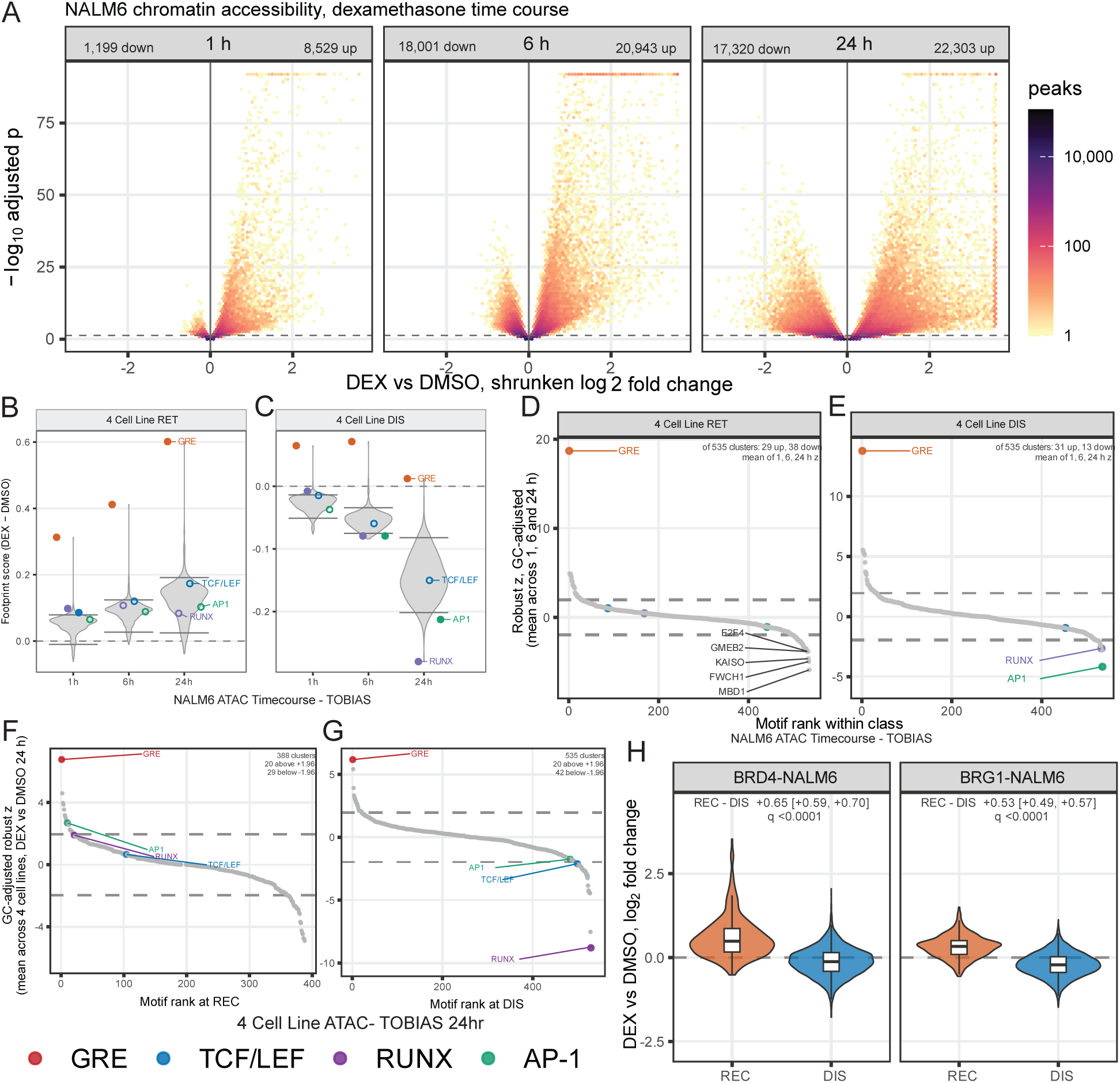
Genome-wide chromatin accessibility and regulatory responses across GR-LEF1 site classes. **(A)** Genome-wide changes in chromatin accessibility in NALM6 cells following 1, 6, and 24 h DEX treatment relative to matched DMSO controls. Volcano plots show differential accessibility across the consensus ATAC-seq peak set. **(B-C)** TOBIAS differential footprinting at GR+LEF1 (B) retention (RET) and (C) dissociation (DIS) sites across the ATAC-seq time course. Violin plots show the distribution of DEX-induced differential footprint scores across all evaluated motifs; points indicate representative GRE, TCF/LEF, RUNX, and AP-1 motifs. **(D-E)** Ranking of DEX-induced differential footprint changes across 535 TOBIAS motif clusters at **(D)** RET and **(E)** DIS sites. All 535 clusters met the ≥100 motif-instance inclusion threshold in both classes, compared with 388 clusters meeting this threshold at REC sites (Figure 4D). Clusters were ranked by robust z scores summarizing differential footprinting across the three DEX time points. Dashed lines indicate z= ±1.96. The GRE cluster occupied the highest-ranked position in both classes. **(F-G)** Ranked DEX-associated footprint changes across motif clusters at GR+LEF1 REC (F) and DIS (G) sites using 24-h ATAC-seq data from four B-ALL cell lines. TOBIAS BINDetect footprint changes were summarized across cell lines and ranked within each site class. GRE, RUNX, TCF/LEF, and AP-1 clusters are highlighted. Dashed lines indicate robust z= ±1.96. **(H)** DEX-induced changes in BRD4 and BRG1 occupancy at REC and DIS sites in NALM6 cells. Violin plots show per-site log_2_ fold changes in CUT&RUN signal following DEX treatment relative to DMSO controls.

**Figure S4.**
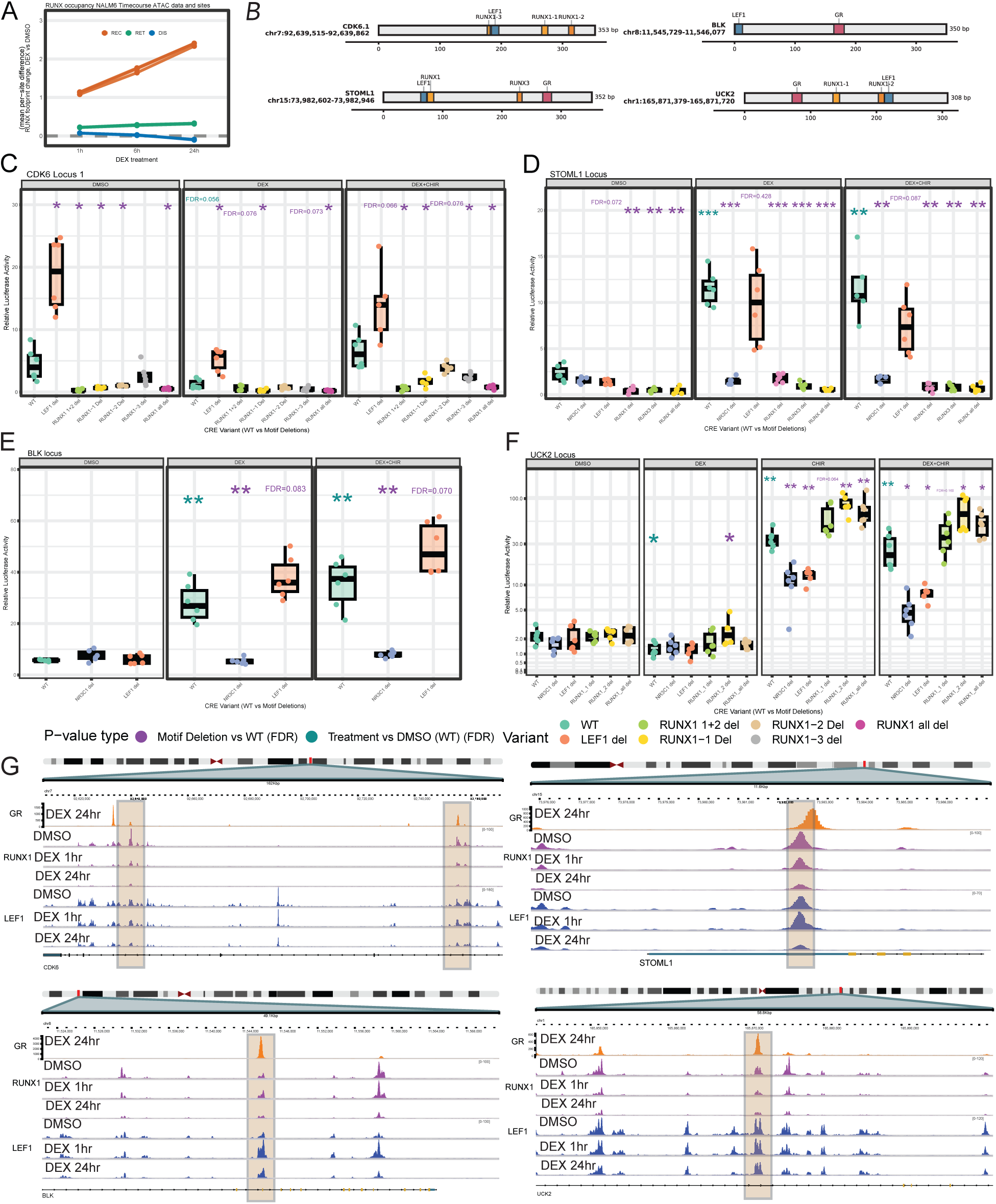
RUNX footprinting and motif-dependent regulatory activity at GR-associated elements. **(A)** Tobias footprinting data showing the change in RUNX family TF footprint score over time in NALM6 cells. **(B)** Diagrams of loci and targeted motif deletions used in luciferase assays below in C-F. **(C-F)** Luciferase activity of GR+LEF1 regulatory elements at CDK6.1 **(C)**, STOML1 **(D)**, BLK **(E)**, and UCK2 **(F)** containing WT sequence or targeted deletions of LEF1, GR, and/or RUNX1 motifs, as indicated. Reporter activity was measured following DMSO, DEX, CHIR, or DEX+CHIR treatment as shown for each element. **(G)** Representative genome-browser tracks showing GR, RUNX1, and LEF1 occupancy at the regulatory elements tested by luciferase assay under the indicated treatment conditions. For panels C-F, motif-deletion constructs were compared with the matched WT element within each treatment, and treatment effects were compared with DMSO-treated WT. FDR-adjusted significance is denoted *q < 0.05, **q < 0.01, and ***q < 0.001; exact FDR values are shown where indicated.

**Figure S5.**
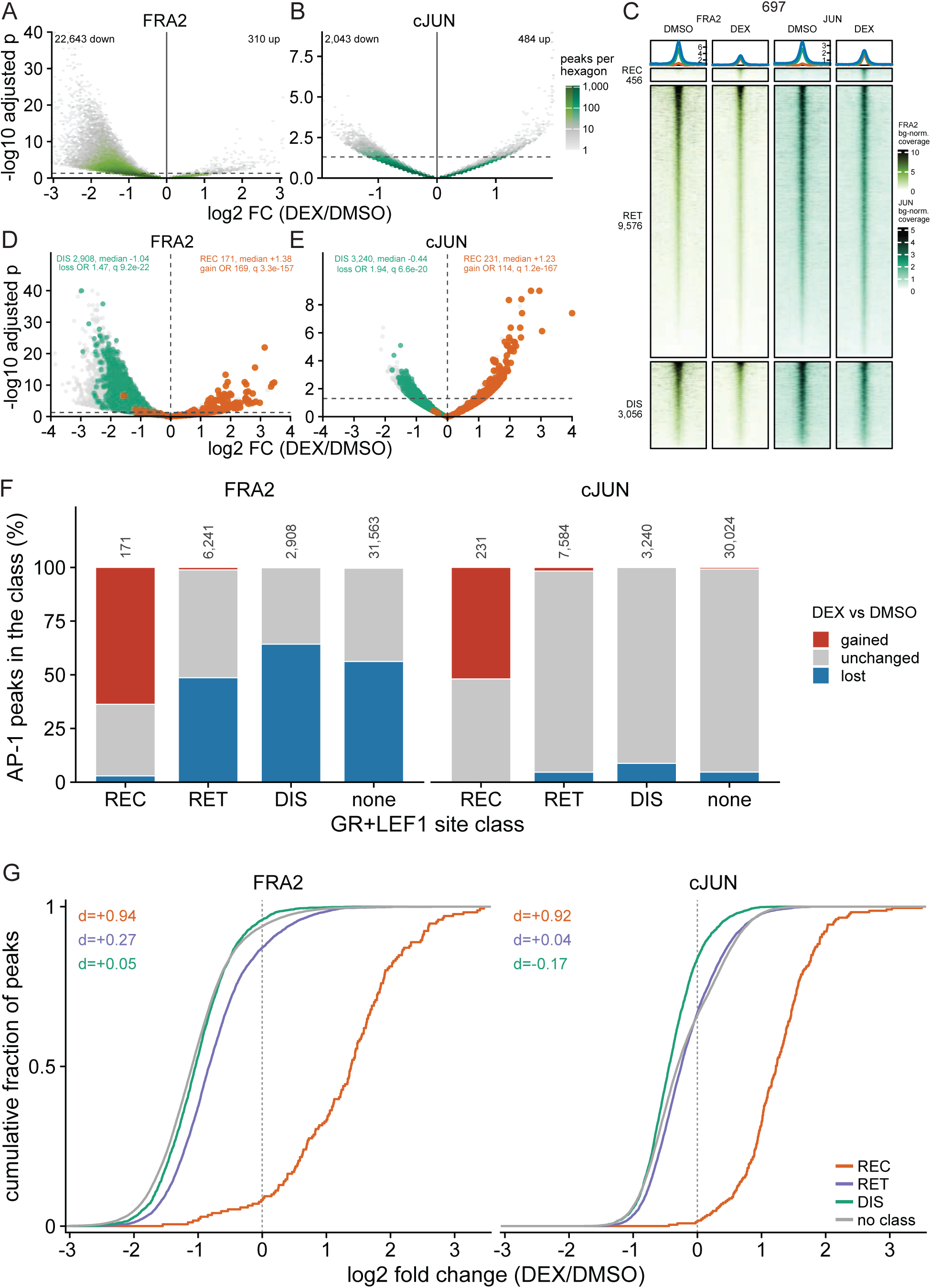
AP-1 transcription factors exhibit class-dependent redistribution following GR activation. **(A-B)** Volcano plots of genome-wide differential FOSL2/FRA2 (A) and JUN/cJUN (B) occupancy in 697 cells following 24 h DEX treatment relative to DMSO. **(C)** Heatmaps and aggregate profiles of FOSL2/FRA2 and JUN occupancy at LEF1-defined GR+LEF1 REC, RET, and DIS sites in 697 cells following DMSO or 24 h DEX treatment. **(D-E)** Differential FOSL2/FRA2 **(D)** and JUN **(E)** occupancy at AP-1 peaks overlapping GR+LEF1 REC or DIS sites. REC-overlapping peaks are shown in green and DIS-overlapping peaks in orange. Among the displayed class-overlapping peaks, REC sites show positive median DEX-associated changes for both factors, whereas DIS sites show negative median changes. **(F)** Fraction of FOSL2/FRA2 and JUN peaks overlapping each GR+LEF1 class that significantly increased, significantly decreased, or were not significantly changed after 24 h DEX treatment. **(G)** Cumulative distributions of shrunken DEX-versus-DMSO log₂ fold changes for FOSL2/FRA2 and JUN peaks overlapping REC, RET, or DIS sites, compared with AP-1 peaks that do not overlap a GR+LEF1 site.

**Figure S6.**
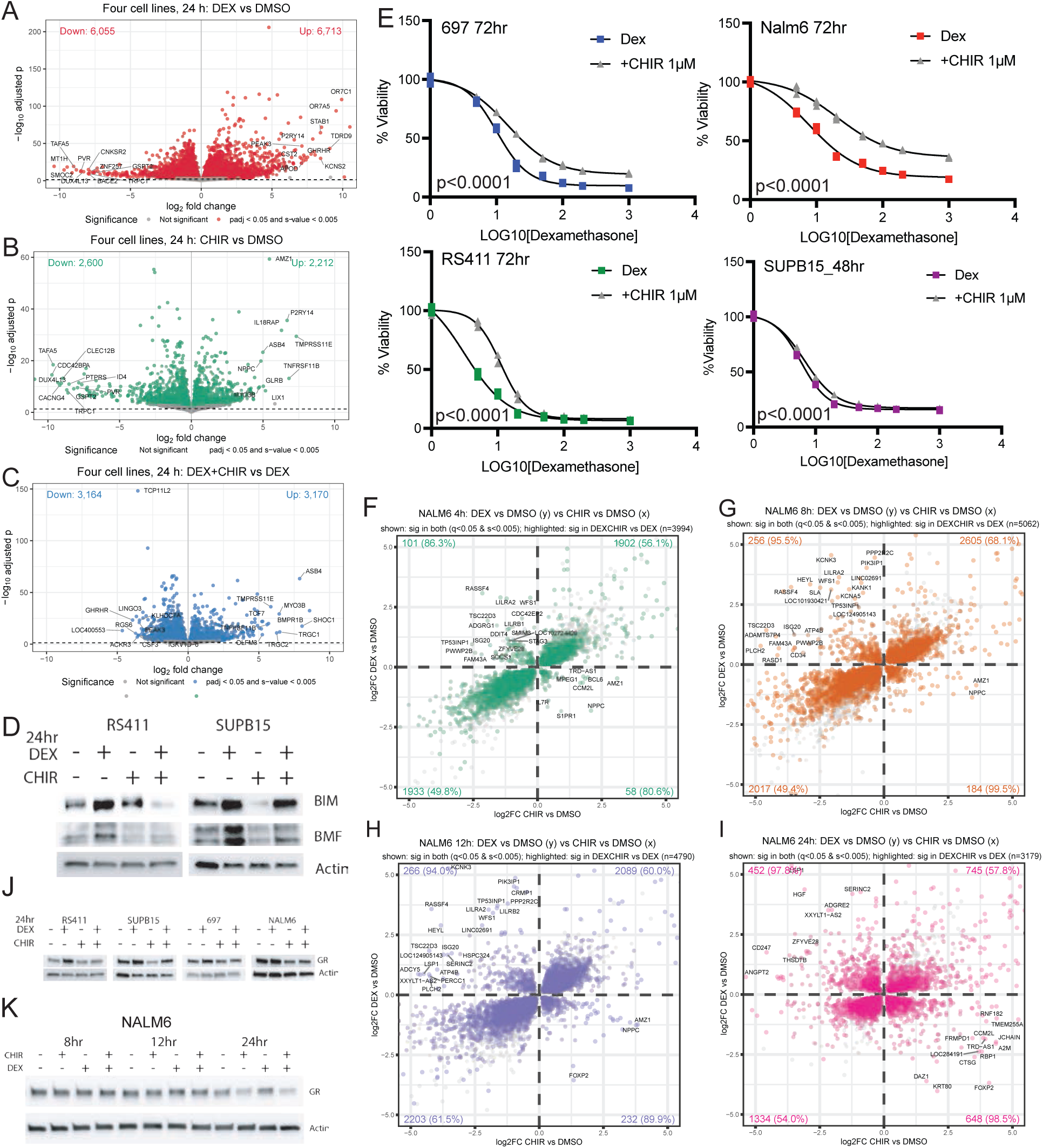
Wnt activation opposes glucocorticoid-responsive gene expression and sensitivity. **(A-C)** Volcano plots of differential gene expression across the four-cell-line panel after 24 h treatment: DEX versus DMSO **(A)**, CHIR versus DMSO **(B)**, and DEX+CHIR versus DEX **(C)**. Genes were considered significantly changed at adjusted p < 0.05 and s-value < 0.005, without a fold-change threshold. DEX altered 12,768 genes (6,713 increased; 6,055 decreased), CHIR altered 4,812 genes (2,212 increased; 2,600 decreased), and addition of CHIR to DEX altered 6,334 genes relative to DEX alone (3,170 increased; 3,164 decreased). **(D)** Immunoblot analysis of BIM and BMF following 24 h DEX, CHIR, or DEX+CHIR treatment in RS411 and SUPB15 cells. **(E)** DEX dose-response viability curves in the presence or absence of 1 µM CHIR in 697, NALM6, RS411, and SUPB15 cells. Addition of CHIR significantly shifted the dose-response curve in all four cell lines (extra sum-of-squares F test, p < 0.0001 for each line) and increased the fitted DEX IC50 from 10.17 to 14.98 nM in 697, 7.83 to 20.83 nM in NALM6, 3.17 to 11.17 nM in RS411, and 5.82 to 6.63 nM in SUPB15. Each treatment arm was normalized to its corresponding vehicle condition. 697, NALM6, and RS411 were assessed at 72 h and SUPB15 at 48 h. **(F-I)** Comparison of log_2_ fold change of DEX and CHIR transcriptional responses in NALM6 cells after 4 h **(F)**, 8 h **(G)**, 12 h **(H)**, and 24 h **(I)**. Genes significantly altered by DEX+CHIR relative to DEX are highlighted. Enrichment of combination-responsive genes in opposing DEX/CHIR quadrants increased over time, with odds ratios of 0.73 at 4 h, 1.27 at 8 h, 1.12 at 12 h, and 2.08 at 24 h (Fisher’s exact test, p = 1.6e-14, 3.6e-10, 1.7e-3, and 1.8e-120, respectively). **(J)** GR protein abundance following 24 h DMSO, DEX, CHIR, or DEX+CHIR treatment across 697, NALM6, RS411, and SUPB15 cells. β-actin serves as a loading control. **(K)** GR protein abundance during an 8-24 h DEX, CHIR, and DEX+CHIR time course in NALM6 cells. GR loss was not apparent until after 12 h of treatment. β-actin serves as a loading control.

**Figure S7.**
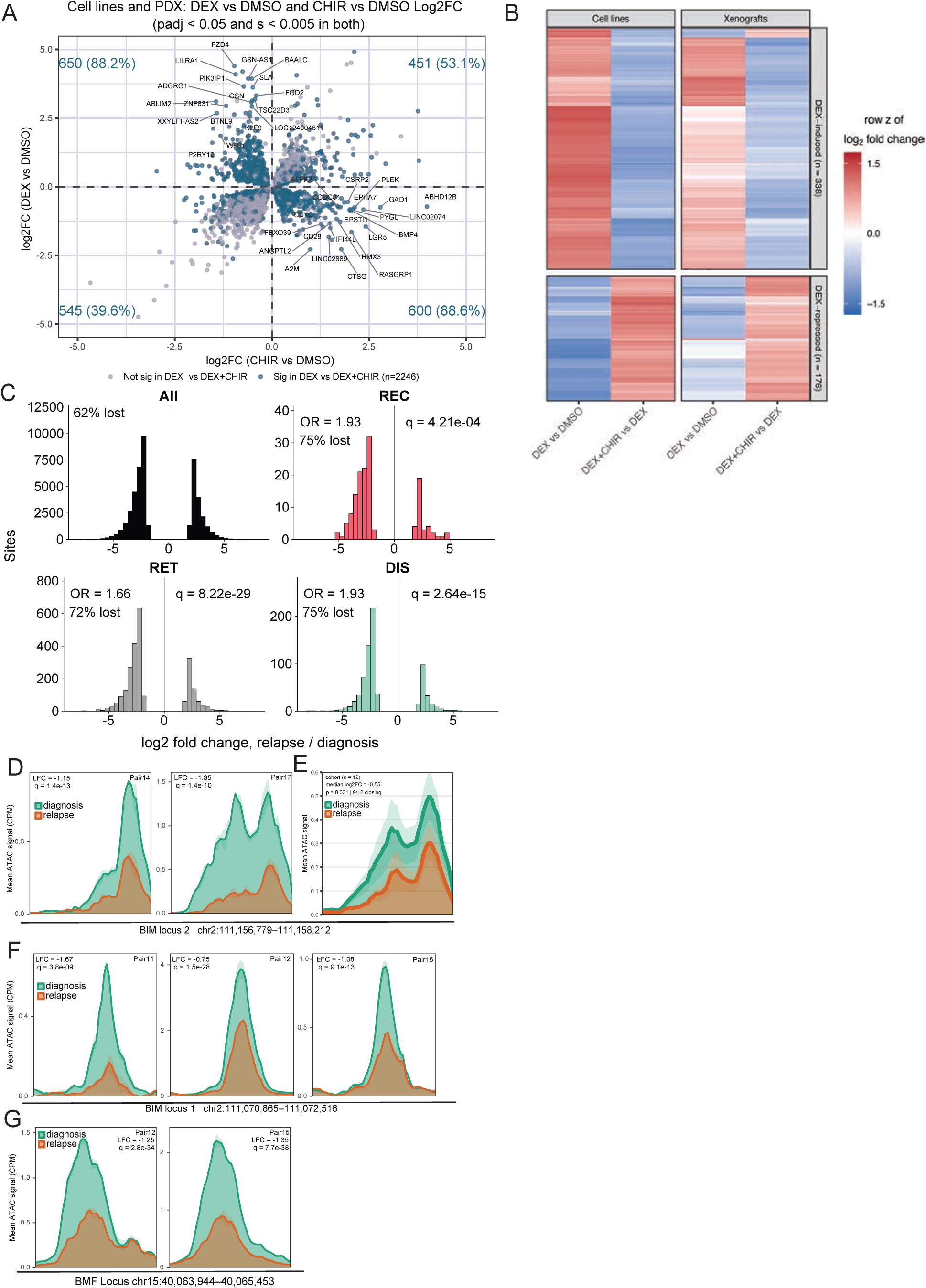
Glucocorticoid-Wnt transcriptional antagonism and diagnosis-relapse accessibility changes in B-ALL. **(A)** Comparison of DEX- and CHIR-induced transcriptional responses across four B-ALL cell lines and five B-ALL patient-derived xenografts (PDXs). Genes significantly regulated by both treatments are shown; highlighted genes are additionally altered by DEX+CHIR relative to DEX. Quadrant enrichment was assessed by Fisher’s exact test. **(B)** Heatmap of genes exhibiting GC-Wnt antagonism in both B-ALL cell lines and PDXs. Shown are log2 fold changes for DEX versus DMSO and DEX+CHIR versus DEX. Rows are scaled by gene and hierarchically clustered. Enrichment of shared antagonized genes was assessed by Fisher’s exact test. **(C)** Distribution of ATAC-seq log₂ fold changes (relapse versus diagnosis) among all strongly remodeled sites (| log₂FC|>2), and within each GR+LEF1 class. Odds ratios, percentages of sites showing accessibility loss at relapse, and significance values are shown. Each class was compared with non-GR+LEF1 sites using a two-sided Fisher’s exact test with Benjamini-Hochberg correction across the three comparisons. Odds ratios above 1 indicate a stronger bias toward loss of accessibility at relapse than in the non-GR+LEF1 comparator: REC, odds ratio = 1.93, q = 4.21e-4; RET, odds ratio = 1.66, q = 8.22e-29; and DIS, odds ratio = 1.93, q = 2.64e-15. **(D)** ATAC-seq coverage at the BIM locus 2 enhancer in two additional diagnosis-relapse pairs showing significant decreases at relapse (DiffBind q<0.05) below the fold-change cutoff (absolute log₂ fold change <2): Pair 14 (log₂ fold change = −1.15; q = 1.4e-13) and Pair 17 (−1.35; q = 1.4e-10). Log₂ fold changes are expressed as relapse relative to diagnosis; lines represent mean accessibility and shading indicates SEM. **(E)** Mean ATAC-seq coverage at the BIM locus 2 enhancer across all 12 diagnosis-relapse pairs, without selection by statistical significance or fold-change magnitude. Accessibility decreased at relapse in 9 of 12 pairs (median log₂ fold change = −0.55; P = 0.031, Wilcoxon signed-rank test of per-pair log₂ fold changes against zero). Lines represent mean accessibility and shading indicates SEM. **(F)** ATAC-seq coverage at the BIM locus 1 enhancer in three diagnosis-relapse pairs showing significant decreases at relapse (DiffBind q<0.05) below the fold-change cutoff used to define strong remodeling (absolute log₂ fold change <2): Pair 11 (log₂ fold change = −1.67; q = 3.8e-9), Pair 12 (−0.75; q = 1.5e-28) and Pair 15 (−1.08; q = 9.1e-13). Log₂ fold changes are expressed as relapse relative to diagnosis; lines represent mean accessibility and shading indicates SEM. **(G)** ATAC-seq coverage at the BMF regulatory element in two diagnosis-relapse pairs showing significant decreases at relapse (DiffBind q<0.05) below the fold-change cutoff (absolute log₂ fold change <2): Pair 12 (log₂ fold change = −1.25; q = 2.8e-34) and Pair 15 (−1.35; q = 7.7e-38). Log₂ fold changes are expressed as relapse relative to diagnosis; lines represent mean accessibility and shading indicates SEM.

